# CRISPR-SID: identifying EZH2 as a druggable target for desmoid tumors via *in vivo* dependency mapping

**DOI:** 10.1101/595769

**Authors:** Thomas Naert, Dieter Tulkens, Tom Van Nieuwenhuysen, Joanna Przybyl, Suzan Demuynck, Matt van de Rijn, Mushriq Al Jazrawe, Benjamin Alman, Paul J. Coucke, Kim De Leeneer, Christian Vanhove, Savvas N. Savvides, David Creytens, Kris Vleminckx

## Abstract

Cancer precision medicine implies identification of tumor-specific vulnerabilities associated with defined oncogenic pathways. Desmoid tumors are soft-tissue neoplasms strictly driven by Wnt signaling network hyperactivation. Despite this clearly defined genetic etiology and the strict and unique implication of the Wnt/β-catenin pathway, no specific molecular targets for these tumors have been identified. To address this caveat, we developed fast and semi-high throughput genetic *Xenopus tropicalis* desmoid tumor models to identify and characterize novel drug targets. We used multiplexed CRISPR/Cas9 genome editing in these models to simultaneously target a tumor suppressor gene (*apc*) and candidate dependency genes. Our methodology CRISPR/Cas9 Selection mediated Identification of Dependencies (CRISPR-SID) uses calculated deviations between experimentally observed gene editing outcomes and deep-learning-predicted double strand break repair patterns, to identify genes under negative selection during tumorigenesis. This revealed *EZH2* and *SUZ12*, both encoding polycomb repressive complex 2 components, and the transcription factor *CREB3L1*, as genetic dependencies for desmoid tumors. *In vivo* EZH2 inhibition by Tazemetostat induced partial regression of established autochthonous tumors. *In vitro* models of patient desmoid tumor cells revealed a direct effect of Tazemetostat on Wnt pathway activity. CRISPR-SID represents a potent novel approach for *in vivo* mapping of tumor vulnerabilities and drug target identification.

**Significance Statement:** CRISPR-SID was established in the diploid frog *Xenopus tropicalis* for *in vivo* elucidation of cancer cell vulnerabilities. CRISPR-SID uses deep learning predictions and binomial theory to identify genes under positive or negative selection during autochthonous tumor development. Using CRISPR-SID in a genetic model for desmoid tumors, treatment-recalcitrant mesenchymal tumors driven by hyper-activation of the Wnt signaling pathway, we identified *EZH2* and *SUZ12*, both encoding critical components of the polycomb repressive complex 2, as dependency genes for desmoid. Finally, we demonstrate the promise of EZH2 inhibition as a novel therapeutic strategy for desmoid tumors. With the simplicity of CRISPR sgRNA multiplexing in *Xenopus* embryos the CRISPR-SID method may be applicable to reveal vulnerabilities in other tumor types.

## Introduction

The identification of novel cancer vulnerabilities is paramount to enable therapeutic discoveries and to improve patient prospects. Recent technological advances allow genetic dependency mapping across a wide array of cancer cell lines at an unprecedented speed (Dempster et al., 2019). At the same time, the introduction of TALEN and CRISPR/Cas has empowered straightforward genome engineering in a large number of animal models previously recalcitrant to genetic studies. A limiting experimental factor for mammalian organisms is the large-scale introduction of the genome-editing reagents in early embryos. However, for aquatic vertebrates such as fish and amphibians with external development and large-size embryos, simple micro-injections allow fast, straightforward and multiplexed introduction of these genome editing compounds. Over the past years, we harnessed TALENs and CRISPR/Cas9 in the true diploid model *Xenopus tropicalis* empowering this amphibian species as a cost-efficient platform for cancer biology (Naert et al., 2016; Naert et al., 2020a; Van Nieuwenhuysen et al., 2015). We further proposed the use of CRISPR/Cas9 to validate cancer cell vulnerabilities *in vivo* by inactivating suspected genetic dependencies in developing tumors (Naert and Vleminckx, 2018). Clonal cancer growth is an inherently Darwinian process, where genetic mutations that increase fitness are under positive selection and mutations disfavoring fitness or growth are under negative selection pressure (Maresch et al., 2016; Weber et al., 2015). By monitoring the selection for specific patterns of CRISPR/Cas9 editing outcomes during autochthonous tumor development, we here establish an experimental pipeline for *in vivo* elucidation of cancer cell vulnerabilities. In brief, we use deep learning to forecast CRISPR/Cas9 gene editing outcomes and subsequently employ binomial statistics to ascertain deviations between expected and observed editing patterns as a measure of tumoral selection mechanisms, thereby identifying genetic vulnerabilities.

Desmoid tumors are orphan-type soft-tissue neoplasms driven strictly by Wnt signaling network hyperactivation (Alman et al., 1997; Tejpar et al., 1999). Desmoid tumors can occur either sporadic and harbor activating missense mutations in the *CTNNB1* gene, or can be associated with germline loss-of-function mutations in the tumor suppressor gene *APC* (associated with familial adenomatous polyposis (FAP) or Gardner’s syndrome). Desmoid tumors do not metastasize and pose a low risk of death in sporadic disease but they confer substantial complications and, dependent on the location of the tumor, cause severe morbidity or lead to amputation of one of the extremities. However, in FAP patients, which normally undergo colectomy, desmoid tumors are the leading cause of morbidity and mortality (Nugent et al., 1993). Despite the clearly defined genetic etiology and the strict and unique implication of the Wnt/β-catenin pathway, no systemic specific molecular therapy for the tumors exists. A number of agents such as the tyrosine kinase inhibitor Sorafenib and the gamma-secretase inhibitor Nirogacestat have been shown to exert activity against desmoid tumors (Gounder et al., 2018; Kummar et al., 2017). However, in both cases the actual drug target and the underlying molecular mode of action in the desmoid tumor cells remains unknown (Gounder, 2015).

Here, using multiplexed CRISPR/Cas9 based genome editing, we performed desmoid tumor genetic dependency mapping directly *in vivo* using *Xenopus tropicalis*. We show novel genetic dependencies during autochthonous desmoid tumorigenesis and report pre-clinical data revealing Ezh2 as a druggable vulnerability in desmoid tumors.

## Results

### Wnt signaling network modulation initiates desmoid tumorigenesis in *Xenopus tropicalis*

Previously, we reported penetrant induction of desmoid tumors in F0 *X. tropicalis* by TALENs-mediated genome engineering of the tumor suppressor gene *apc* (Van Nieuwenhuysen et al., 2015). From these, we generated a stable F1 *X. tropicalis* line containing a germline heterozygous one base pair deletion in the mutation cluster region (MCR) of the *apc* gene (c.4120del, *apc*^MCR-Δ1/+^) (Table S1A). In phenocopy to Gardner syndrome patients, *apc*^MCR-Δ1/+^ animals develop desmoid tumors, showing 100% penetrance (*n*=7) when monitored at the age of five years (Fig. 1A). Desmoid tumors present on MRI as collagen-rich neoplasms, appreciable by T2-weighted hyperintensity, and exhibit conventional key histopathological hallmarks of long sweeping fascicles composed of uniform and slender fibroblasts/myofibroblasts (Fig. 1A) (Zreik and Fritchie, 2016). In the meantime, we similarly induced desmoid tumors by CRISPR/Cas9-mediated targeting of *apc*, which further facilitates downstream multiplexed gene inactivation studies. For this, we similarly targeted the mutation cluster region of *apc* with CRISPR/Cas9 and performed genome editing in (Table S1B) (Tran et al., 2010). Upon injection of 8-cell stage embryos carrying a Wnt/β-catenin reporter construct in the two dorsal-vegetal blastomeres, which contribute primarily to the anterior endoderm and dorsal mesoderm, subsequent post-metamorphic *X. tropicalis* presented with fully penetrant development of multiple desmoid tumors (*n*=10) at three months of age (Fig. 1B). Epifluorescence exposed active Wnt signaling, and histopathology demonstrated the presence of all relevant conventional hallmarks of desmoid tumors (Fig. 1B). Deep PCR amplicon sequencing of a desmoid tumor revealed the presence of biallelic *apc* frameshifting mutations in the MCR, encoded by the final exon, with as a predicted consequence expression of a truncated Apc protein (Table S1C). Finally, desmoid tumors could also be induced by CRISPR-base editing (BE3) mediating a targeted C-to-T conversion in the *ctnnb1* gene (Fig. 1C) (Komor et al., 2016; Shi et al., 2019). As such, we modulate the conserved S45 phosphorylation site of β-catenin in *X. tropicalis*, recapitulating a clinically relevant human S45F missense mutation (Fig. 1C) (Penel et al., 2017). The S45 residue is normally phosphorylated by the casein kinase 1, which primes the β-catenin protein for subsequent phosphorylation by the glycogen synthase kinase 3β in the destruction complex, and thereby earmarks it for ubiquitination and degradation by the proteasome. The stabilized S45F mutant β-catenin protein drove desmoid tumorigenesis in 87.5% of F0 animals (*n=*8) via monoclonal expansion of C-to-T edited cells at three months of age (Fig. 1C). Collectively, we describe three novel genetic models for desmoid tumors in *X. tropicalis* by modulating the Wnt signaling pathway, which can now be exploited for genetic dependency mapping and pre-clinical drug studies.

**Figure 1.**
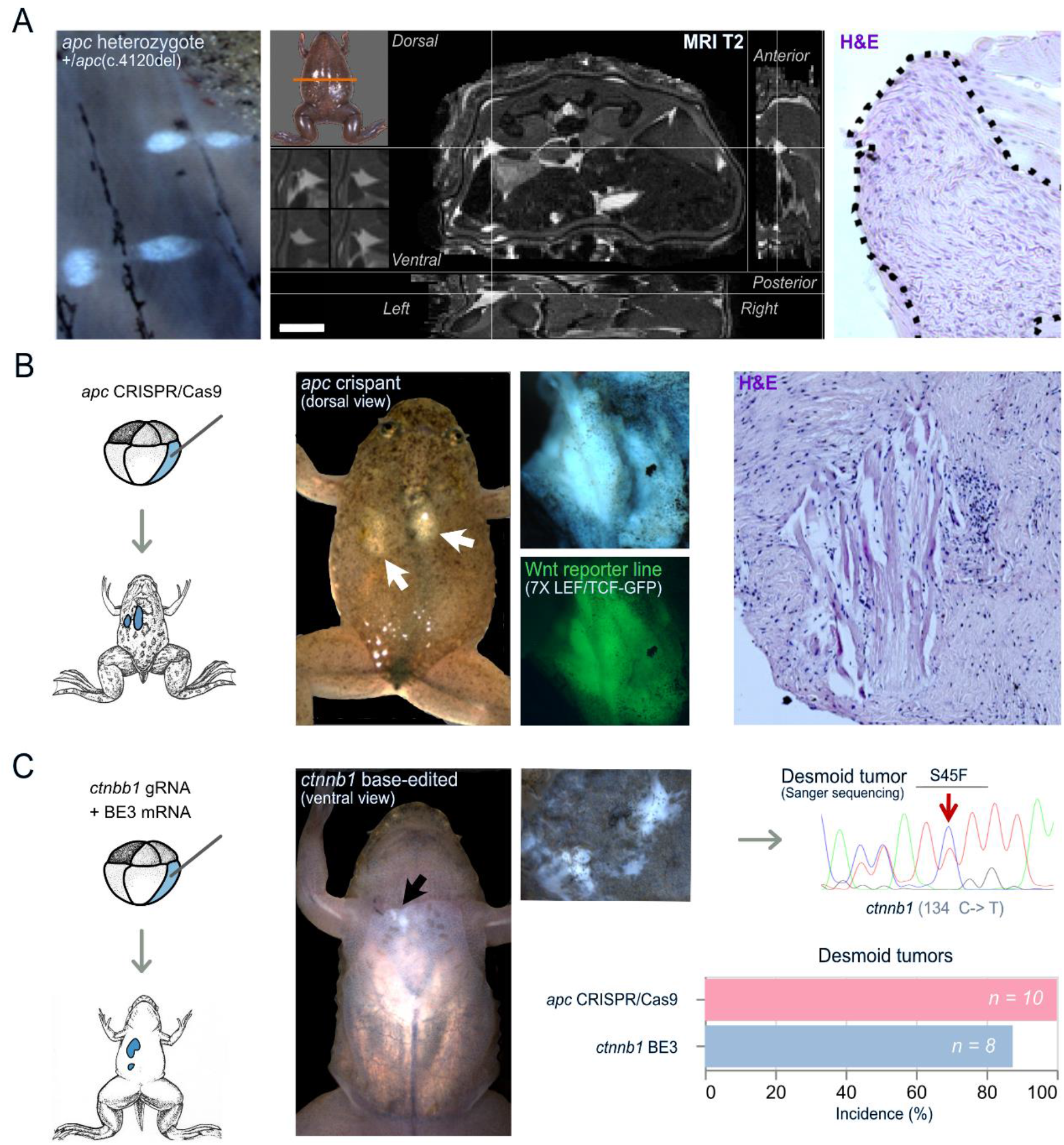
Hyperactivation of the Wnt/β-catenin signaling pathway via truncation of Apc or constitutive activation of β-catenin induces desmoid tumors in *X. tropicalis*. **(A)** *Xenopus tropicalis* carrying a germline heterozygous one base pair deletion in the mutation cluster region (MCR) of the *apc* gene (c.4120del, *apc*^MCR-Δ1/+^) develop desmoid tumors with a 100% penetrance at three years of age (left panel). The desmoid tumors present as T2 hyperintense foci on MRI (middle panel, intersection of the crosshairs) and are characterized by classical desmoid tumor histopathology of long sweeping fascicles composed of uniform and slender fibroblasts/myofibroblasts (right panel). **(B)** *apc* CRISPR/Cas9 injections in the vegetal-dorsal blastomere leads to mosaic mutant animals that manifest several desmoid tumors. These desmoid tumors exhibit active and high Wnt signaling activity as demonstrated by GFP expression using a Wnt reporter line (*WntREs:dEGFP*) (Tran et al., 2010) (middle panel). Classical histopathology of desmoid tumors with long sweeping fascicles of bland fibroblasts/myofibroblasts exhibiting, in this case, local invasion into musculature (right panel). **(C)** Injection of a BE3 cytosine base editor with a *ctnnb1* gRNA leads to development of desmoid tumors. Sanger sequencing of a dissected desmoid tumor demonstrates base pair edits culminating in amino acid changes corresponding the human S45F mutation in *CTNNB1* associated with sporadic desmoid tumorigenesis. Penetrance of desmoid tumors by three months of age is 100% for *apc* CRISPR/Cas9 injected animals and 87.5% for *ctnnb1* base-edited animals.

### CRISPR-SID: quantitative assessments of positive and negative selection pressure on gene inactivation during tumorigenesis

To further explore the use of our novel CRISPR/Cas9-mediated *apc* desmoid tumor model for genetic dependency mapping, we first selected suspected genetic dependencies. We re-analyzed available transcriptomics studies and identified genes which are consistently upregulated in desmoid tumors compared to other fibrotic lesions (Table S2) (Guo et al., 2013; West et al., 2005). In addition, we integrated protein druggability, via DGIdb (Cotto et al., 2018), and mined published desmoid tumor literature (Table S3). As such, we generated a shortlist of ten suspected genetic dependencies in desmoid tumors: *hmmr*, *adam12*, *fap-α*, *mdk*, *nuak1*, *lox*, *pycr1*, *creb3l1*, *pclaf* and *ezh2* (Chen et al., 2014; Colombo et al., 2011; Jung et al., 2013; Misemer et al., 2014; Reversade et al., 2009; Skubitz and Skubitz, 2004; Tolg et al., 2003). For these suspected dependencies, we validated high-efficiency gRNAs by injection in *X. tropicalis* embryos followed by deep amplicon sequencing of the targeted regions (Table S1D).

Next, we developed a novel methodology, CRISPR-SID (**<u>CRISPR</u>**/Cas9 **<u>S</u>**election mediated **<u>I</u>**dentification of **<u>D</u>**ependencies), to ascertain selection pressure towards specific CRISPR/Cas9-induced mutational spectra enriched during autochthonous genetic tumor development. This selection pressure can manifest itself during repair of the CRISPR/Cas9 induced double strand break as positive, via selection for biallelic frameshift mutations in tumor suppressors, or as negative, via counterselection of biallelic frameshift mutations in genetic dependencies. This results from Darwinian natural selection mechanisms occurring *in vivo* within the mosaic CRISPR/Cas9-mutated *Xenopus* during tumor initiation and development. We wanted to quantitatively assess these selection mechanisms at play, by comparing CRISPR/Cas9 editing outcomes in a previously reported machine-learning prediction algorithm, and via deep amplicon sequencing of *Xenopus* tissues under different degrees of selection pressure.

We first deployed the machine learning algorithm InDelphi, which accurately predicts CRISPR/Cas9 editing outcomes. While InDelphi was trained in human cell lines and in mouse embryonic stem cells (Shen et al., 2018), we recently reported that the algorithm is also highly predictive for double strand break repair outcomes in *Xenopus* and zebrafish embryos (Naert et al., 2020b). The model provides us with high-accuracy prediction of genome editing outcomes for our gRNAs, in the absence of tumoral Darwinian selection mechanisms. We co-injected gRNAs targeting each suspected genetic dependency together with *apc* gRNA, and Cas9 protein in blastomeres contributing to the anterior endoderm and dorsal mesoderm in *X. tropicalis* (Table S1E). Post-metamorphosis, each developing desmoid tumor was subjected to deep amplicon sequencing and its mutational spectrum at each CRISPR/Cas9 target site was classified (Fig. 2A-B) (Table S4). We next employed binomial statistics to ascertain whether the classification of the mutation spectrum occurring in desmoid tumors is binomially likely or unlikely given the InDelphi deep-learning predictions (Supplemental Data - Extended Information). A significant deviation in the tumor mutation spectrum, compared to InDelphi model predictions, is indicative for the presence of Darwinian selection mechanisms that either favor or disfavor inactivation of specific genes during desmoid tumor development.

**Figure 2.**
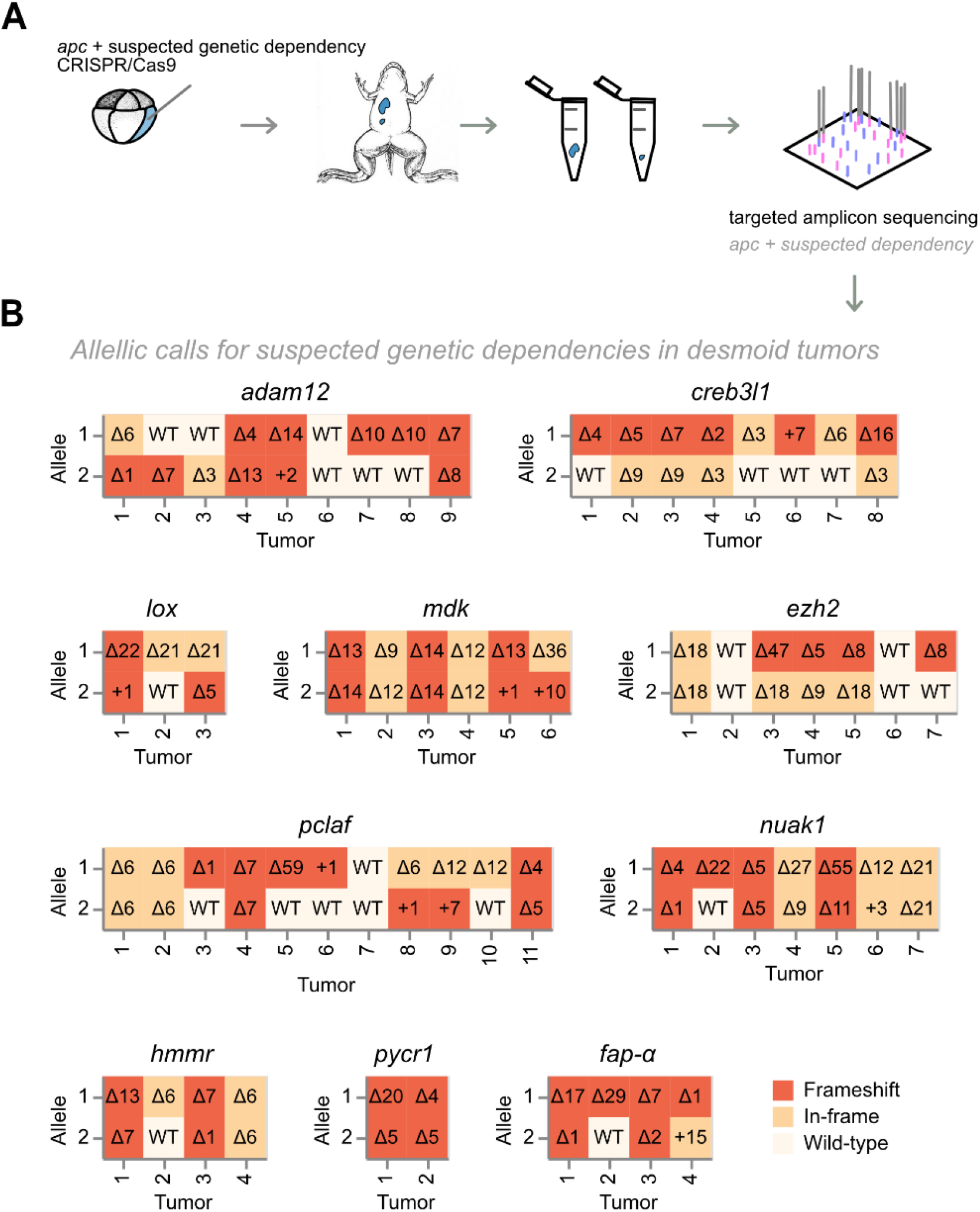
Assessing and classifying editing outcomes at CRISPR/Cas9 target sites of candidate dependency genes in *apc* mutant desmoid tumors. **(A)** *Xenopus tropicalis* embryos are co-targeted at *apc* and respectively one suspected genetic dependency (*e.g. adam12*). Desmoid tumors were dissected from three month old post-metamorphic animals and both CRISPR/Cas9 target sites were subjected to targeted amplicon sequencing to determine gene editing outcomes. **(B)** Indels and allelic status at the suspected dependency CRISPR/Cas9 target sites in dissected desmoid tumors. Frameshift mutations (FS) are indicated in dark-red, in-frame (IF) mutations in ochre, while wild-type allelic calls are indicated in off-white. Numbers in boxes show specific INDEL mutations, where Δn indicates a deletion of n nucleotides and +n indicates insertions of n nucleotides. For eight suspected genetic dependencies, desmoid tumors were retrieved with biallelic frameshift mutations (demarcated by boxed numbers). In contrast, for *ezh2* and *creb3l1*, biallelic frameshift mutations were never retrieved.

### CRISPR-SID ascertains positive selection for biallelic *apc* frameshift mutations in desmoid tumors

Illustrating CRISPR-SID by example, classifying the mutational spectrum of *apc* in desmoid tumors, we observed a high-rate of biallelic *apc* frameshift mutations (Supplemental Fig. S1A). Using InDelphi CRISPR/Cas9 gene editing outcome predictions for biallelic frameshift mutations in a given desmoid tumor (*p*=0.81) (Supplemental Fig. S1B), we employed binomial statistics to calculate the probability that 100% of the tumors would carry biallelic frameshifts in absence of selection (*x*=61, *n*=61). This probability is extremely low and indicative of positive selection pressure for biallelic *apc* mutations in desmoid tumors (P[X=61]<0.01) (Supplemental Fig. S1B). Using CRISPR-SID we thus certify the known role of *apc* as a tumor suppressor in desmoid tumor biology, based solely on deviations between tumor-sampled and InDelphi-predicted *apc* editing outcomes. Evidently, this experimental observation is in direct agreement with the well-described biology of desmoid tumors and the role of *apc* as a tumor suppressor gene (Alman et al., 1997; Latchford et al., 2007).

To validate, we used the same rationale to compare the *apc* gene editing outcomes sampled from whole early embryos and analyzed by deep amplicon sequencing, to *apc* gene editing outcomes in the desmoid tumors. In line with InDelphi model predictions, binomial statistics revealed the probability of the desmoid tumors to have this specific classification of *apc* gene editing outcomes as unlikely to occur by random chance (P[X=61]<0.01) (Supplemental Fig. S1B).

### CRISPR-SID ascertains negative selection pressure and identifies *ezh2* and *creb3l1* as genetic dependencies for desmoid tumors

Classifying the mutation spectrum of each of the ten suspected genetic dependencies (Fig. 2A), we observed for eight genes no significant deviations between InDelphi CRISPR/Cas9 editing outcome predictions and the experimental editing outcomes obtained in the desmoid tumors (Fig. 2A-B, Fig. S2). Importantly, for this statistical analysis only the tumors in which both alleles of the targeted dependency are mutated, are taken into account. As such, this analysis *in se* is independent of cutting efficiency of a particular gRNA. As an example, using InDelphi predictions of the probability on *adam12* biallelic frameshift mutations in a given desmoid tumor (*p*=0.58), we employed binomial statistics to calculate the chance that, as observed, 75% of those tumors that were edited in both alleles, would harbor biallelic frameshift mutations (*x*=3, *n*=4). This probability is high (P[X=3]>0.05) and indicative of lack of any selection pressure for a specific *adam12* CRISPR/Cas9 editing outcome (Fig. 2A, Fig 3A). Biologically, the lack of deviation implies that, for these eight genes, desmoid tumors can develop with biallelic inactivation of said genes, excluding them as genetic dependencies for desmoid tumorigenesis.

**Figure 3.**
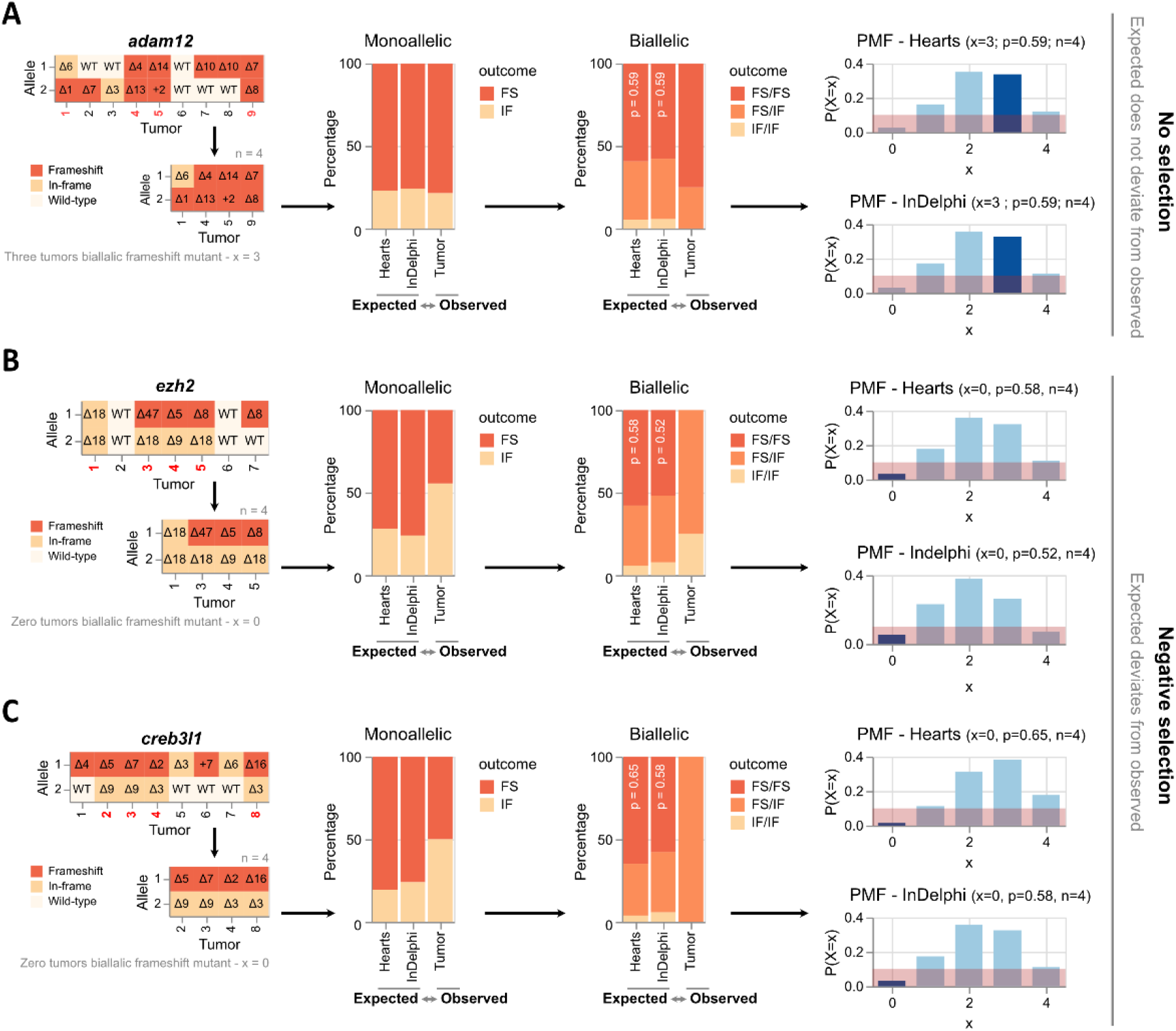
Statistical probability of *ezh2* or *creb3l1* biallelic frameshift mutations in desmoid tumors in relation to deep-learning predictions of gene editing outcomes. **(A)** Lack of selection mechanisms towards specific *adam12* gene editing outcomes during desmoid tumorigenesis. The CRISPR/Cas9 editing outcomes for *adam12* in desmoid tumors containing bi-allelic mutations (in-frame + frameshift), are likely and do not deviate from what is predicted by InDelphi and experimentally sampled in heart tissue. Given the *adam12* editing outcomes sampled in hearts, *i.e*. in absence of selective pressure, the probability of a single biallelic mutant desmoid tumor to have biallelic frameshift editing outcomes is 59%. The probability of sampling 60% (3/5 tumors) with biallelic frameshift *adam12* allelic status is likely (probability is 35%). Similarly, using InDelphi predictions the probability of sampling this proportion of desmoid tumors with *adam12* biallelic frameshift mutations is likely given binomial theory (probability is 34%). Red demarcation represents a 10% probability interval. PMF, probability mass function. **(B-C)** Editing outcomes for *ezh2* and *creb3l1* remain similar between InDelphi predictions and experimental observations in the heart of *X. tropicalis*. Significant deviations from the two former values are seen in the gene editing outcomes occurring in desmoid tumors. In line, the probabilities of never sampling biallelic frameshift mutations in *ezh2* and *creb3l1* within desmoid tumors is binomially unlikely (probability is 5% or smaller) given the editing outcomes in absence of tumor selection (heart tissue and InDelphi).

In direct contrast, we observed for two suspected dependencies, *ezh2* and *creb3l1*, significant deviations between the InDelphi CRISPR/Cas9 editing outcome predictions and the actual experimental mutation patterns obtained by targeted deep amplicon sequencing of sampled desmoid tumors. For both genes, only tumors were obtained where at least one allele of the dependency gene was either wild type or had an in-frame INDEL mutation. For *ezh2* the probability that 0% of those tumors that have both alleles mutated, have in fact no biallelic frameshift mutations (*x*=0, *n*=4) is unlikely (P[X=0]<0.05) given InDelphi predicted biallelic editing outcome probabilities (*p*=0.58) (Fig. 3B). Similarly, the probability that 0% of the tumors would have biallelic *creb3l1* frameshift mutations is unlikely (P[X=0)<0.05) (*x*=0, *n*=4, *p*=0.52) (Fig. 3C). We ruled out biases due to aberrant InDelphi predictions by determining the CRISPR/Cas9 editing outcomes in the hearts of animals from this respective injection setup. As the heart receives major contribution from the injected blastomeres, the editing outcome patterns for *ezh2* and *creb3l1*, in their respective setup, can also be used to determine editing outcomes in absence of tumoral selection pressure. Given these, the chance that 0% of the tumors in the *ezh2* (Fig. 3B) and *creb3l1* (Fig. 3C) setups develop without biallelic frameshift mutations is considered unlikely (both P[X=0]<0.05) (Table S6A).

Taken together, our CRISPR-SID *in vivo* dependency mapping revealed that genetic disruption of *lox*, *adam-12*, *mdk*, *hmmr*, *nuak1*, *fap-α*, *pclaf* and *pycr1* does not prevent desmoid tumor development. In contrast, we observed significant deviations from the expected CRISPR/Cas9 editing outcomes in tumors for the *ezh2* and *creb3l1* genes. This negative selection indicates that *ezh2* and *creb3l1* are genetic dependencies for desmoid tumors.

### Desmoid tumor development critically depends on the activity of the polycomb repressive complex 2

EZH2 functions as the catalytic subunit in the polycomb repressive complex 2 (PRC2), mediating epigenetic alterations via its C-terminal histone methyl transferase SET domain (Kim and Roberts, 2016). EZH2 inhibition is rapidly gaining traction as a therapeutic strategy and an EZH2 inhibitor, Tazemetostat (Tazverik), has recently been FDA-approved via orphan drug designation for treatment of refractory follicular lymphoma and for epithelioid sarcoma, a rare soft tissue cancer (Girard et al., 2021; Gounder et al., 2020; Hoy, 2020). Given the well-tolerated clinical profile of Tazemetostat (Duan et al., 2020) we set-out to validate EZH2 as a dependency in desmoid tumors. Encouragingly, we observed profound H3K27me3 immunoreactivity in *Xenopus* desmoid tumors indicative of high PRC2 activity, in line with human desmoid tumor biopsies (Supplemental Fig. S3A) (Mito et al., 2017).

We further validated that the *ezh2* editing outcomes in tumors are indicative for negative selection towards *ezh2* biallelic frameshift mutations. As outlined above, we found that, in line with *in vitro* CRISPR/Cas9 screens (Shi et al., 2015), selection pressure for functional Ezh2 protein manifests itself via a higher than expected rate of in-frame mutations. Inherent to the selection model, this evidently imposes the condition that the observed *ezh2* variants indeed encode functional proteins. Exploiting the conservation of sequence identity between human, *Anolis carolinensis* and *X. tropicalis* EZH2 protein (Brooun et al., 2016) (Supplemental Fig. S4), we generated a structural scaffold to map EZH2 protein variants retrieved from desmoid tumor deep amplicon sequencing (Fig. 4A). These three in-frame deletion protein variants map within a loop region at the periphery of the EZH2 subunit, which we determine likely to be structurally tolerated (Fig. 4B). Further, we found that these protein variants express *in vivo* using *Xenopus* embryo expression assays and retained both expression and H3K27 trimethyltransferase function upon overexpression in HEK293T cells (Fig. 4C). This illustrates, next to negative selection against biallelic *ezh2* frameshift mutations in tumors, a tolerance for in-frame variants encoding mutant but functional Ezh2 protein. In brief, under strong selection pressure due to CRISPR/Cas9 editing at the *ezh2* target locus, we show that desmoid tumors can escape imposed selection pressure by harboring functional in-frame *ezh2* mutations.

**Figure 4.**
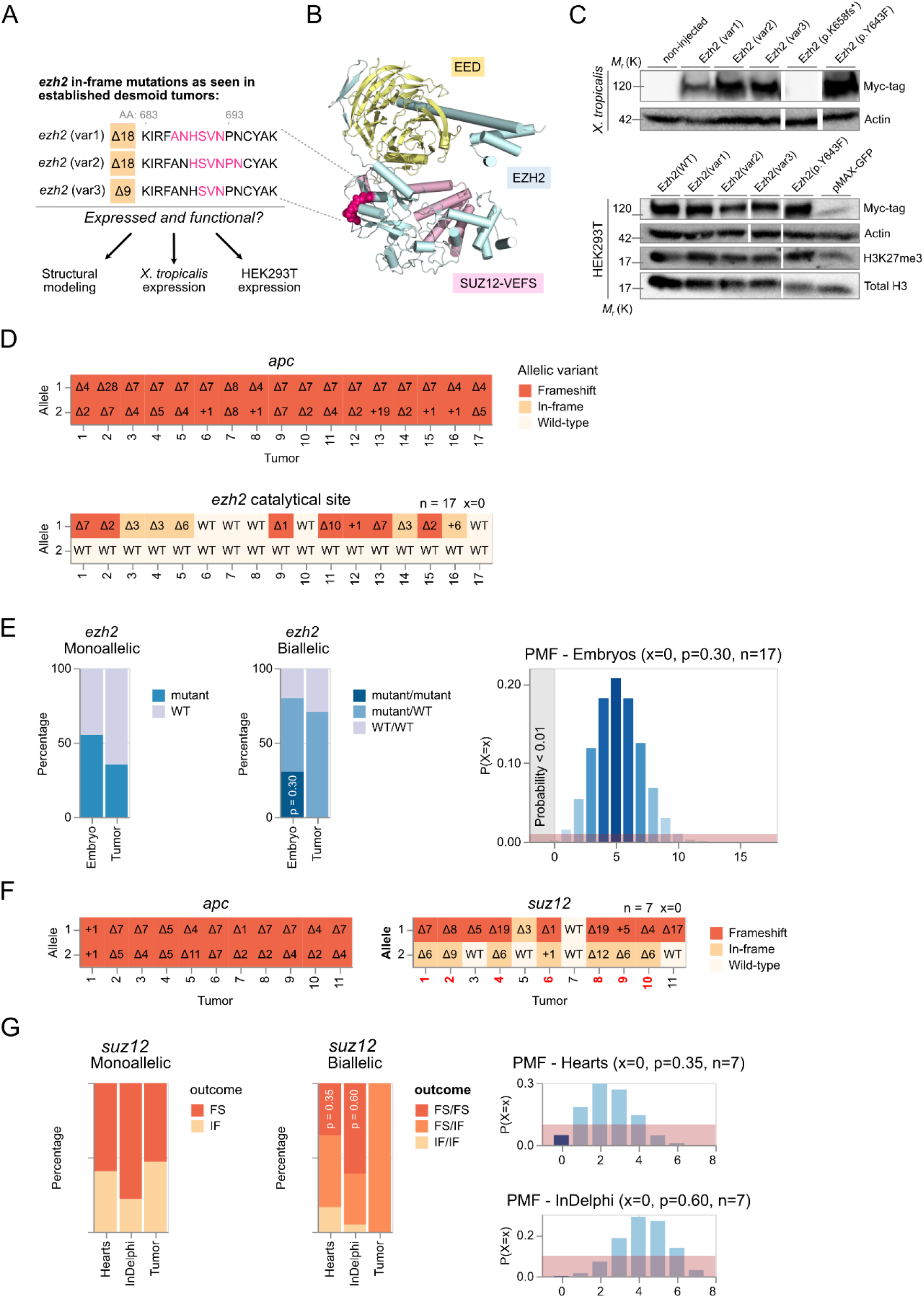
Assessing the role of the polycomb repressive complex 2 in desmoid tumorigenesis. **(A)** In-frame *ezh2* variants observed in desmoid tumors (See Fig. 3B). **(B)** Mapping of identified deletion mutations in the EZH2 subunit of the polycomb repressive complex 2 (PCR2). The deletion variants occur in a loop (magenta) at the periphery of the Ezh2 subunit and are expected to be well tolerated structurally. **(C)** Expression studies of Ezh2 protein variants in *X. tropicalis* and HEK293T cells. EZH2(p.Y643F) (corresponds to EZH2(p.Y642F) in human) and EZH2(p.K658fs*) variants function as positive and negative control respectively. Actin and Total H3 are shown as loading controls. Myc-tagged Ezh2 protein variants express in *X. tropicalis* **(Top)** and in HEK293T cells (**Bottom**) and remain functional as they increase the levels of H3K27me3, when compared to the negative control (pMAX-GFP). Immunoblots were cut and pasted (white spaces) for clarity. Unaltered original blots are shown in Supplementary Fig. S7. **(D)** INDELs and allelic status at the suspected dependency CRISPR/Cas9 target sites in desmoid tumors. Frameshift mutations (FS) are indicated in dark orange, in-frame mutations (IF) in ochre and wild-type (WT) in off-white. **(E)** Gene editing outcomes at the *ezh2* exon encoding the Ezh2 catalytical pocket reveals selection towards retention of at least one wild-type *ezh2* allele. Since each mutation (frameshift or in-frame) at this location can be considered as loss-of-function (LOF), we calculate the probability of a single desmoid tumor to have biallelic LOF *ezh2* editing to be 30%, given *ezh2* editing efficiencies in embryos from the same injection setup. Therefore, the probability of sampling 0% (17 tumors) without biallelic LOF allelic status is very unlikely (probability < 0.01%) according to binomial theory. Red demarcation represents a 1% probability interval. **(F-G)** The *suz12* CRISPR/Cas9 editing outcomes, sampled in desmoid tumors, are unlikely and deviate from the expected. Given the *suz12* editing outcomes sampled in hearts, the probability of a single biallelic mutant desmoid tumor to have biallelic frameshift editing outcomes is 35%. The probability of sampling 0% (0/7 tumors) with biallelic frameshift *suz12* allelic status is unlikely (probability is <5%). Similarly, using InDelphi predictions, the probability of sampling this proportion of desmoid tumors with *suz12* biallelic frameshift mutations is unlikely given binomial theory (probability is <0.01%). This demonstrates selection mechanisms towards specific *suz12* gene editing outcomes during desmoid tumorigenesis. Red demarcation represents a 10% probability interval.

Previous studies have demonstrated that negative selection pressure in CRISPR screens increases dramatically when targeting specific protein domains driving the dependency since small in-frame deletions are in most cases not tolerated in critical functional domains (Michlits et al., 2020; Shi et al., 2015). This, as each such mutation, whether in-frame or frameshift, can be considered a loss-of-function mutation. To investigate the importance of the histone methyltransferase activity of Ezh2 in its function as a dependency factor, we designed a gRNA targeting the SET-domain at valine 659 (*ezh2^catalytical site^*). With *ezh2^catalytical site^*, we observed strong selection pressure on retaining at least one wild-type non-edited allele in all desmoid tumors sampled (Fig. 4D). Based on editing efficiencies of the *ezh2^catalytical site^* gRNA in embryos from the same injection setup, the chance of recovering seventeen tumors with none showing biallelic *ezh2* editing events is binomially highly unlikely (*p*<0.01) (Fig. 4E) (Table S6B). This confirms that the methyltransferase activity of Ezh2 is indeed crucial towards its activity as a dependency factor for desmoid tumor formation.

Importantly, there is growing evidence that histone methyltransferases like EZH2 can have methyltransferase-dependent functions not associated with PRC2 (Wang and Wang, 2020). Hence, we investigated whether *suz12*, encoding an essential structural subunit of PRC2, could be similarly identified as a genetic dependency for desmoid tumor development. For this, we designed a gRNA targeting *suz12* at asparagine 338 (*suz12^N338^*). We performed CRISPR-SID revealing negative selection on biallelic *suz12* frameshift mutations using *a priori* experimental data in the heart (*p=*0.048; n=7) and InDelphi predictions (*p*<0.01; n=7) (Fig 4F-G).

Collectively, these experiments demonstrate that both *ezh2* and *suz12* are genetic dependencies during desmoid tumor development with selection pressure specifically on the SET-domain of the Ezh2 protein. This makes a strong case for a dependency on functional PCR2, mediating H3K27Me3 marks, during desmoid tumorigenesis.

### Chemical inhibition of Ezh2, via Tazemetostat, elicits a therapeutic response in established desmoid tumors

To investigate whether the genetic dependency of desmoid tumorigenesis for PRC2 could be harnessed towards a molecular therapy, we investigated whether EZH2 inhibition could elicit a therapeutic response in animals with established desmoid tumors. To exploit our *apc* mosaic mutant CRISPR/Cas9 *X. tropicalis* desmoid tumor model for pre-clinical drug studies, we exposed desmoid-tumor bearing froglets via the rearing water to the EZH2-inhibitor Tazemetostat (Tazverik/EZP-6438) (Fig. 5A). A critical issue when administrating a compound via the rearing water of the animals, is to ascertain active absorption and biodistribution. Using a dose of 10μM Tazemetostat we demonstrated a reduction, but not complete elimination, of H3K27me3 levels in the livers of drug-exposed animals, indicative of moderate *in vivo* Ezh2 inhibition (Fig. 5B). With this concentration, no adverse effects were detected on the well-being of the exposed animals. A three-week Tazemetostat exposure resulted in a reduction or stasis in desmoid tumor size, whereas sustained tumor growth was observed in the control arm exposed to 0.1% DMSO (*p*<0.05) (Fig. 5C-D). While the externally visible desmoid tumors normally had a cloudy milky appearance with smooth borders, after Tazemetostat treatment they appeared more nodular and compacted, with irregular margins (Fig. 5C). Hence, we assessed the collagen content and organization via picrosirius red staining and polarization microscopy (Fig. 5E). This revealed that Tazemetostat-treated desmoid tumors have a significantly higher proportion of collagen organized in fibers, when compared to mock-treated animals (*p*<0.05) (Fig. 5F).

**Figure 5.**
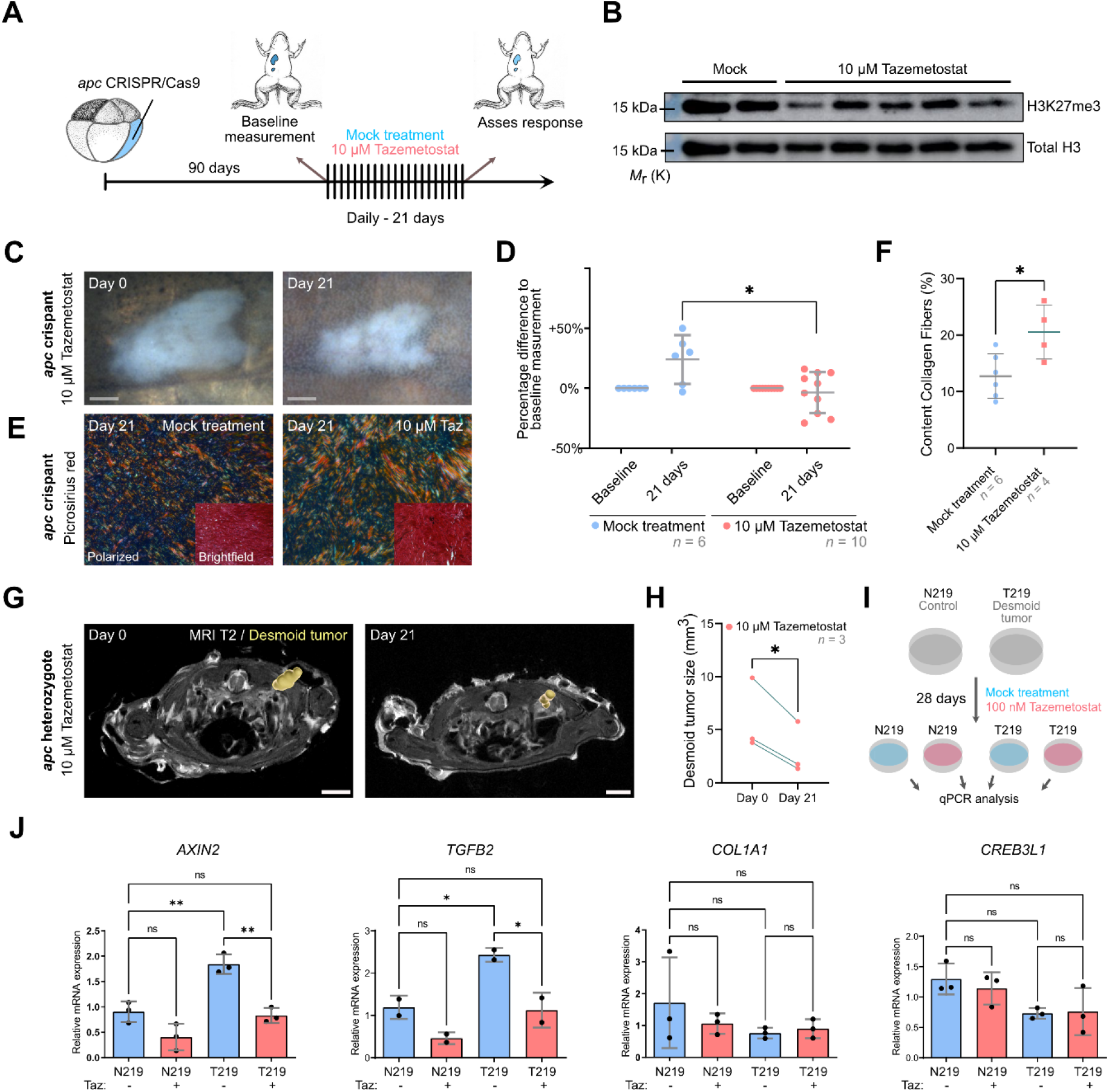
Ezh2 inhibition via Tazemetostat elicits a therapeutic response in desmoid tumors. **(A-B)** Tazemetostat-treated F0 *apc* crispants show a reduction in H3K27me3 levels in the liver (unaltered original scans of blots Supplementary Fig. S7). **(C-D)** Quantitatively, desmoid tumor size reveals stable or progressive disease in the control arm, while in Tazemetostat-exposed animals regression or stasis of tumors could be observed. Each data-point represents one desmoid tumor, note that some animals developed multiple tumors (two-way ANOVA with repeated measurements, Table S6D, \**p* < 0.05). **(E-F)** Picrosirius Red staining with polarized light detection. The degree of alignment of collagen fibers, is significantly increased in Tazemetostat treated tumors when compared to mock treated tumors. (t-test, Table S6D, \**p* < 0.05) **(G-H)** Desmoid tumors in *apc*^MCR-Δ1/+^ heterozygotes respond treatment with Tazemetostat (paired t-test, Table S6D, \**p* < 0.05). **(I)** Treatment scheme of human desmoid tumor cells (T219) and paired normal cells derived from the fascia (N219) with Tazemetostat (100 nM) for four weeks. **(J)** Quantitative RT-PCR analysis of AXIN2, TGFB2, COL1A1 and CREB3L1 expression in DMSO (blue bars) or Tazemetostat (red bars) treated cells. (one-way ANOVA, Table S6D, \**p* < 0.05 \*\**p* < 0.01). Grey scale bar is 1 mm and white scale bar is 3 mm.

While the above mentioned *apc* crispant model is very penetrant and fast, it has the potential caveat, inherent to the method, that the two alleles of the tumor suppressor gene *apc* are already disrupted at an early developmental stage. However, FAP and Gardner’s syndrome patients harbor an inherited heterozygous *APC* mutation and desmoid tumor development occurs upon a second hit in the *APC* gene or upon loss-of-heterozygosity (LOH). Similarly, for sporadic desmoid tumorigenesis somatic mono-allelic activating mutations in the *CTNNB1* gene are acquired at adulthood or juvenile age. In order to investigate whether Tazemetostat could as well affect established desmoid tumors in the context of FAP or Gardner’s syndrome, we first performed magnetic resonance imaging (MRI) on *apc*^MCR-Δ1/+^ animals to detect the possible presence of desmoid tumors. MRI scans of *apc* heterozygotes (aged > 3 years; *n=*3) revealed hyperintense T2 signals indicative of desmoid tumors (Cassidy et al., 2020). These animals were exposed for 4 weeks to Tazemetostat (added at a concentration of 10μM to the rearing water) after which another T2 scan was taken. Dissection and histopathology confirmed the presence of several desmoid tumors, especially in between the dorsal back muscles and on the abdominal wall. Upon necropsy, we retrospectively localized the most prominent desmoid tumors, present between the back muscles, on the MRI scan images. This revealed a significant shrinkage of all three desmoid tumors under observation after Tazemetostat treatment (*p* < 0.05) (Fig. 5G-H and Supplemental Fig. S5).

In sum, the drug treatment experiments show that established desmoid tumors in *Xenopus tropicalis* are dependent on the methyl transferase activity of Ezh2, which can be modulated by Tazemetostat treatment.

### Tazemetostat inhibits the Wnt/β-catenin transcriptional response in human *CTNNB1* mutant primary tumor cells

In order to expand our identified dependency on functional PRC2 in desmoid tumors towards the human situation, we initially confirmed the presence of strong nuclear EZH2 immunoreactivity in human desmoid tumor specimens (Supplemental Fig. S3B-C). In line with this, H3K27me3 immunoreactivity had previously been reported as a histopathological characteristic of desmoid tumors (Mito et al., 2017). Interestingly, tying back to our initial dependency mapping of *creb3l1* in this study, we also found nuclear immunostaining of the N-terminally cleaved fragment of CREB3L1, indicative of an active role as transcription factor in clinical desmoid tumors (Chen et al., 2014) (Supplemental Fig. S3C).

Finally, in order to investigate whether Tazemetostat has an effect on the growth of human desmoid tumor cells cultured *in vitro*, we compared three primary cell lines obtained from clinical desmoid tumors and also included an equal number of normal fibroblasts (Mercier et al., 2018). A dose-response curve with different concentrations of Tazemetostat (5μM - 0.1nM) showed absence of tumor-specific sensitivity for growth inhibition (Supplemental Fig. S6).

Next, in order to investigate a potential effect at the level of Wnt signaling responses, we selected a desmoid tumor cell line (T219) harboring the *CTNNB1*^S45F^ mutation, a mutation frequently found in sporadic desmoid tumors (Trautmann et al., 2020), for which paired normal fibroblasts (N219) were available. The desmoid tumor and normal cells were treated for 4 weeks with 100 nM Tazemetostat. At the end of the treatment, quantitative RT-PCR was performed to evaluate expression of genes highly transcribed in desmoid tumors (*COL1A1*, *COL1A2*, *CREB3L1*, *TGFB2*) and of the *AXIN2* gene, which is a general direct Wnt response gene (Jho et al., 2002; Timbergen et al., 2019). Interestingly, Tazemetostat reduced *AXIN2* and *TGFB2* expression levels in desmoid tumor cells to the level observed in DMSO treated normal cells. The expression of *COL1A1*, *COL1A2* and *CREB3L1* was not affected. Taken together, we conclude that EZH2 inhibition by Tazemetostat has a direct effect downstream or at the level of the β-catenin protein and normalizes Wnt signaling responses in human desmoid tumor cells.

## Discussion

Desmoid tumors are uniquely driven by Wnt signaling hyperactivation and lack current standard of care (Gounder et al., 2018). Here, we describe how novel CRISPR/Cas9-mediated desmoid tumor models in *X. tropicalis* were leveraged to identify Ezh2 as a druggable dependency.

We modulated the Wnt signaling pathway using both classical CRISPR/Cas9 for introducing truncating INDEL mutations in the *apc* gene, thereby mimicking familial adenomatous polyposis, and CRISPR base editing, for establishing an activating missense mutation (S45F) in beta-catenin, associated with sporadic desmoid tumors (Komor et al., 2016; Shi et al., 2019). We also established a stable *X. tropicalis* line (*apc^ΔMCR/+^*) phenocopying presentation of desmoid tumors as seen in FAP and Gardner syndrome (Van Nieuwenhuysen et al., 2015). Importantly, our models all converge on the induction of desmoid tumorigenesis, independent of the specific lesion in the Wnt signaling network. This further enforces the view that desmoid tumors are a consequence of Wnt signaling network hyperactivation in the broad sense (Alman et al., 1997; Tejpar et al., 1999; Trautmann et al., 2020).

Exploiting our novel models we developed a methodology, CRISPR-SID, to identify cancer genetic dependencies. Dependency mapping allows to identify genetic perturbations that negatively affect the potential for either tumor initiation, tumor cell growth or tumor cell survival, as those perturbations will be depleted via Darwinian selection (Boehm et al., 2021). However, ascertaining negative selection in any process is challenging without high confidence knowledge of that process outcome in absence of selection pressure. Therefore, for *in vivo* dependency mapping, which relies on the absence of bi-allelic gene disruption in a candidate dependency gene, one could always argue that biallelic frameshift mutations in this gene could still be observed if a larger sample size would be taken. As, while substantial evidence may prove the absence of biallelic frameshifts in a gene within tumors, there is no direct evidence that denies that this state could exist. This is especially the case for CRISPR/Cas9 experiments where there is never full experimental control on the cutting efficiency of the introduced gRNA, which can be dependent on sequence context and the chromatin status of the target region (Jain et al., 2021; Jensen et al., 2017; Liu et al., 2016).

We believe our current study addresses this conundrum. We provide the framework to ascertain that the lack of sampling biallelic frameshift mutations in a given gene during tumorigenesis is not due to random chance, but rather the result of selection processes. We show that the ever-increasing accuracy by which gene editing outcomes can be predicted by deep convolutional network learning approaches (*a priori*) can be used to demonstrate the deviation from expected gene editing outcomes in processes under selection pressure, such as tumorigenesis (*a posteriori*) (Naert et al., 2020b; Shen et al., 2018). Using this, we unveil selection processes at play during autochthonous desmoid tumorigenesis revealing genetic dependency on *ezh2*, *suz12* and *creb3l1*. This selection pressure can be ascertained by substantial deviations between predicted (x% frameshift DSB repairs) and actual experimentally observed CRISPR/Cas9 editing outcomes (0% frameshift DSB repairs), a fact which we harness to identify cancer cell vulnerabilities. This *in se* turns a negative depletion signal into a positive enrichment signal. As most gRNAs will elicit editing outcomes with a higher frequency of frameshift mutations than in-frame mutations (Shen et al., 2018), the consistent observation of in-frame editing outcomes, due to selection pressure, can yield high statistical certainties with modest sample sizes of tumors. This provides the basis of the CRISPR-SID method. Interestingly, for *in vitro* CRISPR dependency screens, the retention of functional in-frame variants is still considered a nuisance that increases noise within negative selection screens. However, we postulate that, if accurate *a priori* predictions of gene editing outcomes are available for the cell system under scrutiny, selection for in-frame mutants can be harnessed as a positive enrichment signal.

With the potential of our novel CRISPR/Cas9 based *apc* mosaic mutant desmoid tumor model, which combines high penetrance (100%) with short latency (10-12 weeks) of tumor induction, we performed in depth investigation into the nature of the *ezh2* dependency. This is important since EZH2 is best known for its canonical role as a histone methyltransferase in the polycomb repressive complex 2, but has also been documented to show non-canonical functions independent of PRC2, either via methylation of other substrates or even as a transcriptional activator (Wang and Wang, 2020). While in-frame DSB repairs could be observed in the initially targeted CXC domain of ezh2 (Fig. 3B), targeting its catalytic SET domain ascertained manifest selection pressure against either frameshift or in-frame DSB repairs (Fig. 4D-E). Furthermore, we also identified *suz12*, encoding another component of PRC2 (Brooun et al., 2016; Chen et al., 2018), as a dependency gene. This clearly shows a strict dependency on canonical PRC2 activity. In line with our observations, clinical desmoid tumor samples have prominent nuclear localized expression of EZH2 and show strong immunoreactivity for H3K27me3, reflecting high PRC2 catalytic activity (Mito et al., 2017).

Exploiting our *Xenopus tropicalis* model for pre-clinical compound validation, we exposed animals carrying established autochthonous desmoid tumors, either F_0_ *apc* crispants or *apc^ΔMCR/+^* heterozygotes, to the EZH2-inhibitor Tazemetostat. We found consistent reduction or stasis of the desmoid tumors, already after a three- or four-week treatment. Given the fact that Tazemetostat is well tolerated and FDA approved for the treatment of follicular lymphoma and epithelioid sarcoma (Gounder et al., 2020; Hoy, 2020), the drug could hold great promise for repurposing to treat patients with desmoid tumors. Importantly, Tazemetostat treatment in the *Xenopus* model induced a rapid measurable response. This contrasts with the timelines towards clinical responses of the tyrosine kinase inhibitor sorafenib (median time 9.6 months) and the gamma-secretase inhibitor Nirogacestat (> 8 months) currently available to patients afflicted with desmoid tumors (Gounder et al., 2018; Takahashi et al., 2020). Also, for both latter cases the actual drug target involved in the beneficial clinical outcome in desmoid tumors remains unknown. It is also worth noting that both those therapeutic agents elicit no major effects on desmoid tumor cells under laboratory culture conditions, underlining the need to identify novel drugs directly in animal models (Gounder, 2015).

Despite their limitations, such as the absence of a tumor microenvironment, human T219 desmoid tumor cell cultures provided insight into the possible mechanism by which EZH2 inhibition may affect the tumors. We found that Tazemetostat significantly reduced the high expression of the *AXIN2* and *TGFB2* genes to levels observed in control-treated normal fibroblasts. The *AXIN2* gene is a direct and the least tissue-specific transcriptional target of the Wnt/β-catenin pathway (Jho et al., 2002). The *TGFB2* gene is not known as a direct Wnt target but has been found to be upregulated by Wnt stimulation, likely in a tissue-dependent way (Biressi et al., 2014; Bridgewater et al., 2011; Rudolf et al., 2016). As mentioned, we also find the *TGFB2* gene upregulated in human desmoid tumors (Table S2).

Interestingly and importantly, the human T219 desmoid tumor cells carry activating missense mutations in the *CTNNB1* gene that increase the stability of the β-catenin protein by interfering with its casein kinase 1 and glycogen synthase kinase 3β mediated phosphorylation and subsequent ubiquitination (Clevers and Nusse, 2012; Mercier et al., 2018). Hence reduced expression of the *AXIN2* Wnt/β-catenin response gene by Tazemetostat is indicative of a reduced transcriptional response of the β-catenin protein. Since the drug must thereby act either at the level of, or downstream from, the β-catenin protein, this implies that the drug can be effective in the context of both FAP/Gardner syndrome, associated with germline *APC* mutations, and in the context of sporadic desmoid tumors, originating from somatic *CTNNB1* mutations.

At the moment it is still unclear what is the exact nature of the clinical response of the desmoid tumors to the Tazemetostat treatment. One enticing possibility is that the desmoid tumors are addicted to the high signaling activity of the Wnt/β-catenin pathway, which makes them particularly sensitive to the inhibition of the pathway response. However, we failed to detect manifest cell proliferation or cell death responses, neither *in vitro* on human desmoid tumor cells nor *in vivo* in the *Xenopus* desmoid tumors. An alternative mode of action, possibly related to the role of PRC2 in the maintenance of stem cell characteristics, could be that EZH2 inhibition drives the tumor cells into a condition of terminal differentiation. In this context, we found that in the *Xenopus* desmoid tumors, Tazemetostat induced changes in the organization of the collagen fibers, which are excessive in desmoid tumors. Interestingly, it was previously shown that collagen content and cellularity in desmoid tumors are inversely correlated (McCarville et al., 2007). Hence, increased presence of matured collagen fibers may be indicative of a differentiation response in desmoid tumor cells. Intriguingly, CREB3L1, which we also identified as a dependency factor for desmoid tumors, is known as a regulator of collagen synthesis and secretion (Chen et al., 2014; Guillemyn et al., 2019). This may suggest that desmoid tumors critically depend on a deregulated collagen synthesis program, and that modulation of the latter has potential as a therapeutic strategy. A final possibility explaining the activity of Tazemetostat in an *in vivo* context and lack of a tumor-specific response *in vitro*, may point to the involvement of the tumor microenvironment or the immune system, either innate or adaptive, in the drug response (Eich et al., 2020). Experiments addressing these different scenarios are currently ongoing.

This study further empowers the use of *Xenopus tropicalis* for modeling human cancer. A major advantage of the *Xenopus tropicalis* CRISPR-SID model for dependency mapping, is evidently the speed and the straightforwardness of the experimental approach. Micro-injections are technically uncomplicated to perform and multiplexed CRISPR/Cas9 genome editing is well-established (Naert et al., 2020a). Also, since CRISPR-SID is based on positive and negative selection, high efficiency genome editing is not *per se* required. This can be important for candidate dependency genes that are essential for early developmental processes, as in fact is the case for *ezh2* and *suz12* (O’Carroll et al., 2001; Pasini et al., 2004). In this context it is also very convenient that in *Xenopus* early embryos the different blastomeres, contributing to discrete developmental lineages, can be readily distinguished by their pigmentation characteristics and are strictly separated (Moody, 2018). Hence, it is possible to target genome editing to specific lineages and to titrate down gRNA/Cas9 RNP complexes, thereby reducing interference of possible developmental phenotypes (DeLay et al., 2018).

In conclusion, we demonstrate a new experimental approach, CRISPR-SID, to ascertain negative selection in *in vivo* CRISPR/Cas9 experiments using deep learning predictions and binomial theory. We harness this novel approach to identify genetic dependencies on *ezh2*, *suz12* and *creb3l1* during autochthonous desmoid tumorigenesis. Finally, we demonstrate the promise of EZH2 inhibition as a novel therapeutic strategy for desmoid tumors.

## Material and Methods

### Generating the *apc*^MCR-Δ1/+^ line

F0 *apc* mosaics generated by *apc* TALENs injection (Van Nieuwenhuysen et al., 2015) were outcrossed with WT *X. tropicalis*, and offspring was genotyped to identify heterozygotes via heteroduplex mobility assay. One distinct band pattern on heteroduplex mobility analysis (HMA) was determined by deep amplicon sequencing and BATCH-GE (Boel et al., 2016) to be indicative of a one base pair deletion in *apc* and these animals were grouped and further raised for analysis.

### CRISPR/Cas9 genome editing

The *apc* gRNA was designed with the Doench, Hartenian algorithm (Doench et al., 2014). *Mdk*, *wisp-1*, *lox*, *adam-12*, *hmmr*, *nuak1*, *creb3l1*, *FAP-α*, *pclaf^s8^*, *pclaf^G10^*, *pycr1*, *Ezh2^R49^*, *Ezh2^S692^*, *Ezh2^V659^*, *tp53* and *suz12* gRNAs were designed with the CRISPRScan software package (Moreno-Mateos et al., 2015). *In vitro* transcription, purification and quality control of injection-ready gRNA was performed as described before (Table S7A) (Naert et al., 2020b).

### DNA extraction and targeted deep sequencing

In order to analyze CRISPR/Cas9 genome editing efficiencies, embryos, tissue or tumors were incubated overnight at 55 °C in lysis buffer (50 mM Tris pH 8.8, 1 mM EDTA, 0.5% Tween-20, 200 μg/ml proteinase K). For assessing total genome editing efficiency for a specific injection setup we pooled nine stage 46 tadpoles before performing lysis. CRISPR/Cas9-target-site-containing amplicons were obtained by PCR with the primer pairs shown in Table S7B. Deep amplicon sequencing was performed with a validated pipe-line, INDEL frequency data and variant calling for all sequencing was performed with BATCH-GE bio-informatics analysis (Boel et al., 2016).

### Generation of *X. tropicalis* mosaic mutants by CRISPR/Cas9

Wild-type *Xenopus tropicalis* females and males were primed with 20U and 10U PREGNYL^©^ human chorionic gonadotropine (hCG) (Merck), respectively. Natural matings were set-up 2 days later, after boosting the female and male with 150U and 100U of hCG, respectively. Embryos were injected with the concentrations of pre-complexed ribonucleoproteins (RNP) and in the blastomeres described in Table S7C. For all experiments we used NLS-Cas9-NLS (VIB Protein Service Facility, UGent), generated as described before (Naert et al., 2016). All experiments were approved by the Ethical Committee for Animal Experimentation from Ghent University, Faculty of Science and VIB-Site Ghent (License EC2018-079). All methods were carried out in accordance with the relevant guidelines set out by this committee.

### *In silico* differential expression analysis

In order to identify genes that are consistently upregulated in desmoid tumor fibromatosis compared to other fibrotic lesions, we re-analyzed the data from two previously published studies. The first dataset was generated by gene expression microarrays from 8 desmoid tumors, 10 dermatofibrosarcoma protuberans and 13 solitary fibrous tumors (West et al., 2005). The second dataset comprised 7 desmoid tumors compared to 9 different types of fibrous lesions that were profiled by 3SEQ (Cotto et al., 2018; Guo et al., 2013). We performed multiclass differential expression analyses using SAM for microarray data and SAMseq for 3SEQ data. We identified 251 genes (Table S2) that were upregulated in desmoid tumors in both datasets with contrast ≥2 and false detection rate ≤0.01. We performed functional annotation of these 251 genes using Gene Set Enrichment Analysis (GSEA) (MSigDB, Molecular Signatures Database v6.2). Druggability was assessed via the DGIdb database (Cotto et al., 2018).

### CRISPR-SID

For assessment of the gRNA efficiencies for the candidate dependency genes, each gRNA was injected together with Cas9 recombinant protein as pre-complexed RNP, either unilaterally or bilaterally, in two-cell *X. tropicalis* embryos (Table S7C). Genome editing efficiencies were determined as described above. CRISPR-SID was performed by multiplex CRISPR/Cas9, thus generating double crispants for both *apc* and respectively each of the eleven putative dependency genes under scrutiny. Genome editing efficiencies were determined as described above. Animals were further raised until post-metamorphosis. Please note that, considering the fluctuating *apc* editing efficiencies across setups, direct comparison of tumor incidence rates to delineate impact of the candidate dependency gene knockout on desmoid tumorigenesis is impossible. For each animal developing one or multiple desmoid tumors, targeted PCR amplicon sequencing was performed for the *apc* and the respective candidate dependency gene on the desmoid tumors and, in most cases, the heart. Noteworthy, the heart is a tissue receiving significant contribution from the vegetal-dorsal lineage and as such will serve as a proxy to determine the probability of frameshift mutations occurring for a specific gRNA in absence of selection pressure. In practice, each isolated desmoid tumor will have a varying content of non-neoplastic cells, which will be edited at basal efficiencies similar to the vegetal-dorsal lineage of the animal. In order to perform a CRISPR-SID analysis, we require monoclonal tumors and as such all tumors which showed more than two frequent mutation read variants in the *apc* gene were excluded from further analysis. Genes were excluded as genetic dependencies when they demonstrated biallelic frameshift mutations in desmoid tumors. The probability of *y* biallelic frameshifts was calculated using the binomial probability distribution, which takes the form 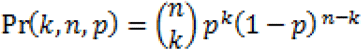. The calculated probability of *k* biallelic frameshift mutations is then function of the number of biallelic frameshifts, *k*, the number of samples, *n*, which is known, and the probability of the occurrence of biallelic frameshifts *p*, which has been determined either experimentally by genetic analysis of the *ezh2^S692^* and *creb3l1^L207^* INDEL patterns in the heart, or as predicted by inDelphi software (Shen et al., 2018). For a more detailed method, see Supplemental Data – Extended Information.

### Imaging, histology and immunohistochemistry

Pictures of desmoid-tumor bearing animals, including the active Wnt signaling within desmoid tumors by dEGFP fluorescence, were taken on a Carl Zeiss StereoLUMAR.V12 stereomicroscope. *X. tropicalis* desmoid tumor tissue samples were harvested from benzocaine euthanized animals, fixed overnight at 4°C in 4% PFA, dehydrated and embedded in paraffin. Tissue sections were cut (5 μm), stained and imaged as described before (Naert et al., 2020a). Clinical human desmoid tumor samples were identified by D.C. and anonymized tissue sections were provided to perform immunofluorescence. Antigen retrieval was performed using a PickCell 2100-Retriever (ProteoGenix) in citrate buffer (10 mM citric acid, 0.1% Tween-20, pH 6). Slides were subsequently blocked with blocking buffer (3% natural serum, 1% BSA, 0.1% Tween-20) and incubated overnight with anti-EZH2 antibody (3748; Abcam), anti-CREB3L1 (AF4080; R&Dsystems) or anti-H3K27me3 (07-449, Merck). Detection was performed with goat anti-rabbit DyLight-594, donkey anti-goat DyLight-594 and goat anti-rabbit DyLight-633, respectively. Counterstaining was performed with Hoechst-33342 and sections were imaged using a Leica TCS LSI zoom confocal microscope. Picrosirius red-polarization detection of collagen fibers in desmoid tumor tissue sections was performed as previously described (Rittie, 2017). Images were acquired on a Axio Observer.Z1 inverted microscope attached to a monochrome Axiocam 506 camera (Carl Zeiss Microscopy GmbH) and analyzed using Fiji software. For quantification of Picrosirius red staining, following image processing was performed. The red channel was isolated from RGB input and masked as a binary. Mask was filtered for elongated collagen fibers based on size exceeding background threshold (>10 pixels) and circularity between 0,4 and 1 (representing elongated polygon blobs). Remaining mask was normalized to the total area of microphotograph containing desmoid tumor and described as percentage content collagen fibers.

### Expression of EZH2 protein variants

C-terminal Myc-epitope-tagged expression plasmid pCS2-MT-EZH2 was generated by cloning *X. tropicalis ezh2* cDNA, excluding the stop codon, in-frame into the BamHI-ClaI sites of the pCS2-MT vector. Specific mutations, NM_00101793.2:c.2163_2180del; NM_001017293.2:c.2165_2182del; NM_001017293.2:c.2171_2179del; NM_001017293.2:c.2068_2074del; NM_001017293.2:c.2127A>T, were introduced in full-length *ezh2* cDNA by the Q5^®^ Site-Directed Mutagenesis Kit (NEB; E0544S). Resulting plasmids were validated by Sanger sequencing, digested with NotI and employed for SP6 *in vitro* transcription of *ezh2* variant mRNA. Two-cell *X. tropicalis* embryos were injected with 500 pg of capped mRNA and allowed to develop to stage 19 and cell extract was prepared in E1A lysis buffer (250 mM NaCl, 50 mM Tris pH 7.4, 0.1% NP-40) containing a complete protease inhibitor cocktail (1:25) (Roche) and centrifuged for 10 min at 14.000 rpm in a microcentrifuge at 4 °C. Cleared supernatant, without yolk contamination, was further used. For transfection experiments, wells were seeded with 200,000 HEK293T cells and transfected the next day using CaCl2 transformation with 100 ng of each of the pCS2-MT-EZH2 plasmids or 100 ng pMAX-GFP plasmid transfection control together with 900 ng of pCS2-MT carrier plasmid. For two replicates, cell extracts were prepared in RIPA buffer (25mM Tris-HCl pH 7.6, 150mM NaCl, 1% NP-40, 1% sodium deoxycholate, 0.1% SDS) containing complete HALT protease and phosphatase inhibitor cocktail (Pierce). For the third replica, cells were harvested in 1x PBS, centrifuged, resuspended and split in two. One part was used for cell extract preparation by RIPA as described before. The other part was used for histone extraction. Histone extracts were prepared as described before (Skrypek et al., 2018), the nucleus was first purified by centrifugation (1X PBS, 0.5% Triton X-100 and 5 mM sodium butyrate) followed by acid extraction (0.4 M HCl) for 1 h at 4 °C and finally the extract was neutralized with 0.5M Na_2_HPO_4_. All protein extracts were denatured in Laemmli buffer, separated by SDS-polyacrylamide gel electrophoresis (20 μg for total protein extracts and 1 μg for histone extracts) and transferred to PVDF. Following antibodies were used for immunodetection: anti-Myc tag (9106; Abcam), anti-β-actin-HRP (47778; Santa Cruz), anti-H3K27me3 (39155; Active-motif) and anti-total H3 (1791; Abcam).

### *In vivo* EZH2 inhibition

Desmoid tumor-bearing *apc* crispants were generated as described before. Ninety-day old animals were randomly assigned to either the treatment or the control arm. Animals were kept in 0.1x MMR containing either 10 μM Tazemetostat (HY-13803; MedChem Express)/0.11% DMSO or 0.11% DMSO. Every day, half of the 0.1x MMR, containing either Tazemetostat or DMSO, was refreshed. A baseline tumor photography was made using a Carl Zeiss StereoLUMAR.V12 stereomicroscope, and a second photograph was taken after 21 days of treatment. Tumor sizes were quantified using ImageJ and statistical analysis was performed using a Two-way ANOVA with repeated measures. For Tazemetostat treatment in the *apd*^MCR-Δ1/+^ line, adult animals were kept in water supplemented with 10 μM Tazemetostat for four weeks, with refreshment of half of Tazemetostat containing water every second day. Before and after the treatment, a whole body MRI scan was taken. For MRI, adult animals were sedated by submersion in 0.2% MS222 (Tricaine) dissolved in water buffered to pH7.0 with Sodium Bicarbonate. Upon disappearance of moving reflexes, animals were briefly submerged in fresh water and wrapped in a wet paper towel and plastic foil. MRI scans were acquired on a 7-Tesla preclinical MRI scanner (Bruker PharmaScan 70/16, Ettlingen, Germany) using a 40 mm quadrature volume transmit/receive radiofrequency coil (Bruker, Ettlingen, Germany). Anesthetized animals were positioned on the scanner’s animal bed. A localizer scan was acquired followed by a T1- and T2-weighted Rapid Acquisition with Relaxation Enhancement sequence to collect anatomical information of thorax and abdomen. The total acquisition time for each scan approximated 9 minutes. After imaging, animals were placed in a tilted container with fresh water, making sure to keep the head in the air. Fresh water was regularly pipetted on the partially submerged animal until breathing and movements became visible after which they were placed back in the original water tanks.

### Cell culture drug treatment and qRT-PCR expression analysis

A primary human desmoid tumor cell line (T219) and a patient matched population of normal fibroblasts (N219) were grown in DMEM supplemented with 10% FCS and seeded in 6-well plates. Cells were treated for four weeks with 100 nM Tazemetostat or with vehicle (0.1% DMSO) with refreshment of half of the medium every second day. Total RNA was isolated from human desmoid tumor cell lines using Trizol reagent (Invitrogen) according to the manufacturer instructions. First-strand cDNA synthesis was performed using the SensiFAST^™^ cDNA Synthesis Kit (Bioline) and RNA concentration levels and purity were determined using a NanoDrop spectrophotometer (Thermo-Scientific). Real-time quantitative PCR was performed using the SensiFAST^™^ SYBR^®^ No-ROX Kit (Bioline) combined with a LightCycler^®^ 480 Real-Time PCR System (Roche). Expression was normalized to housekeeping genes Actb, B2m and Gapd. Data was analysed using the qBase+ software (Biogazelle). All primers including sequences used in this study can be found in Table S7D.

## Author contributions

T.N and K.V designed the study and wrote the manuscript. T.N., D.T., T.V.N. and S.D. were involved in the *Xenopus* experiments. J.P. and M.v.d.R. performed the transcriptome studies in human desmoid tumors. M.A.J. and B.A. derived the human desmoid tumor cells. P.J.C. was involved in the collagen studies and K.D.L. supervised the amplicon deep sequencing. S.N.S. performed the protein structure modeling and C.V. did the MRI analysis. D.C. did histopathological evaluation of the tumor samples.

## Acknowledgements

T.N. was funded by “Kom op tegen Kanker” (Stand up to Cancer), the Flemish cancer society and a PhD fellowship with VLAIO-HERMES (IWT-SB). Research in the authors’ laboratory is supported by the Research Foundation – Flanders (FWO-Vlaanderen) (grants G0A1515N and G029413N), by the Concerted Research Actions from Ghent University (BOF15/GOA/011 and BOF20/GOA/023). Further support was obtained by the Hercules Foundation, Flanders (grant AUGE/11/14), the Desmoid Tumor Research Foundation, the Desmoid Tumour Foundation of Canada and SOS Desmoïde. The authors would like to thank Arne Martens, Nicolas Skrypek and Marja Kreike for helpful suggestions regarding the cell culture experiments. We would like to acknowledge Marjolein Carron and Lana Hellebaut for proof-reading and scientific discussions leading to new relevant insights.

## Notes

The authors declare no conflicts of interest

### Competing Interest Statement

The authors have declared no competing interest.

### Summary of Updates

Additional experiments have been performed and statistical methods have been improved. The revision also contains in vivo MRI imaging of drug responses and analysis of human cell lines.

