## Supplemental Data for "CRISPR-SID: identifying EZH2 as a druggable target for desmoid tumors via *in vivo* dependency mapping"

### Extended Data

#### CRISPR-SID: CRISPR/Cas9 mediated selection of dependencies

These data outline the probabilistic concepts underlying the CRISPR-SID methodology.

**Problem statement**

We provide a mathematical framework for ascertaining negative selection of CRISPR/Cas9 editing outcomes towards null or function-impeded proteins in genes essential for tumorigenesis. Directly ascertaining this negative selection is challenging, as proving the absence of an occurrence is always more challenging that proving the presence of an occurrence. We show the framework to establish burden of proof of negative selection in CRISPR/Cas9 experiments.

**Rationale**

If a tumor critically depends on the expression of protein X, then in no cases should a tumor develop under the absence of the expression of protein X. Introducing CRISPR/Cas9 in this though experiment, this indicates that when targeting the coding sequence for protein X, that biallelic frameshift mutations would never be sampled within single tumors.

**Solution**

**(1)** When a locus is edited, there are two discrete outcome states after repair which take the form:

$$p(Frameshift) = p(FS) = \ probability\ frameshift\ repair$$
$$p(InFrame) = p(IF) = probability\ \ inframe\ repair$$

$$p(FS) + p(IF) = 1$$

**(2)** Under the assumption of biallelic gene editing the following is true and follows a Punnet square logic:

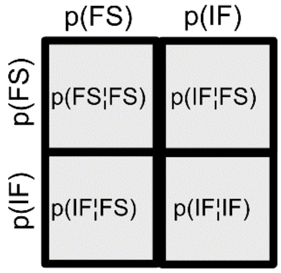


$$p(FS¦FS) + p(IF¦FS)\ + p(FS¦IF)\ + p(IF¦IF) = 1$$

**(3)** CRISPR/Cas9 editing outcomes are probabilistic, with each possible outcome occurring at a rate that is reproducible and predictable. We define:

p(FS) can be determined via two distinct routes

A- InDelphi deep learning predictions

Based on the sequence context surrounding the gRNA cut site the deep learning model predicts the probabilistic percentage gene editing outcome events. This allows for a model predicted chance on monoallelic frameshift mutations p(FS)

B- Experimental observations in tissues edited by CRISPR/Cas9, under the absence of tumoral selection mechanisms

We *(Naert et al, SciRep 2020)*, have previously postulated the following. In *X. tropicalis*, CRISPR/Cas9 reagents are typically injected at an early developmental stage (1 to 16 cell stage). The early embryo rapidly progresses through cell division and as such the delivered CRISPR/Cas9 cuts are repaired individually in the cell population as the embryo grows. will generate a spectrum of mosaic CRISPR/Cas9 mutations events in different cells of the animal. This leads to the following:

**The ratios of cells within the mosaic presenting with certain INDEL variants is representative of the probabilistic outcomes of gene editing towards that specific mutation.**

This assumption was validated using *in vivo* approaches *(Naert et al, SciRep 2020)* and allows us to experimentally obtain the chance on monoallelic frameshift mutations p(FS). For this, we deep amplicon sequence the targeted locus in a pool of CRISPR/Cas9 targeted embryos.

**(4)** The probability for biallelic frameshift mutations [p(FS/FS)] is directly related to the probability for monoallelic frameshifting mutations [p(FS)] derived from (2) and (3) and follows the form:

$$p(FS¦FS) = p(FS)^2$$

**(5)** We can sample the editing outcomes in biallelic edited single desmoid tumors. We can describe these as outcomes from a Bernoulli trial (yes-no question). Either the editing outcomes are biallelic frameshift [$(FS¦FS)$] or they are not [$(IF¦IF)$ or $(IF¦FS)$].

**(6)** Given 1-5, we fit the problem to a traditional binomial distribution which takes the form:

$$f(k,n,p) = \Pr(X = k) = \binom{n}{k} p^k (1-p)^{n-k}$$

This formula described the probability of getting exactly *k* successes, given *n* independent Bernoulli trials given a chance on success of *p*.

Example: Suppose a biased coin comes up heads with probability 0.3 when tossed. The probability of seeing exactly 4 heads in 6 tosses is

$$f(4,6,0.3) = \binom{6}{4} 0.3^4 (1-0.3)^{6-4} = 0.059535$$

However here, we describe the probability of getting exactly *k* tumors without biallelic frameshift mutations, given the chance on biallelic frameshift mutations of *p*(FS¦FS) (derived from (5)) when investigating a total of *n* tumors

Example 1: for *adam12* (Fig. 3A) we derive following:

*p* from InDelphi:

$$p(FS) = 0.77$$
$$p(FS¦FS) = 0.59$$

$$p = 0.59$$

*k* from observations of editing patterns in tumors:

Three tumors carry biallelic frameshift mutations

$$k = 3$$

*n* from the amount of tumors under scrutiny:

Four tumors were investigated

*n = 4*

We derive the probability of these observations:

$$f(k,n,p) = f(3,4,0.59) = 0.3682$$

There is a 36% chance to observe this given InDelphi predictions

Example 2: for *ezh2* (Fig. 3B) we derive following:

*p* from InDelphi:

$$p(FS) = 0.72$$
$$p(FS¦FS) = 0.52$$

$$p = 0.52$$

*k* from observations of editing patterns in tumors:

Zero tumors carry biallelic frameshift mutations

$$k = 0$$

*n* from the number of tumors under scrutiny:

Four tumors were investigated

*n = 4*

We derive the probability of these observations:

$$f(k,n,p) = f(0,4,0.52) = 0.05308$$

There is a mere 5.3% chance to observe this given InDelphi predictions.

### Supplementary Figures

**Supplementary Figure 1:**

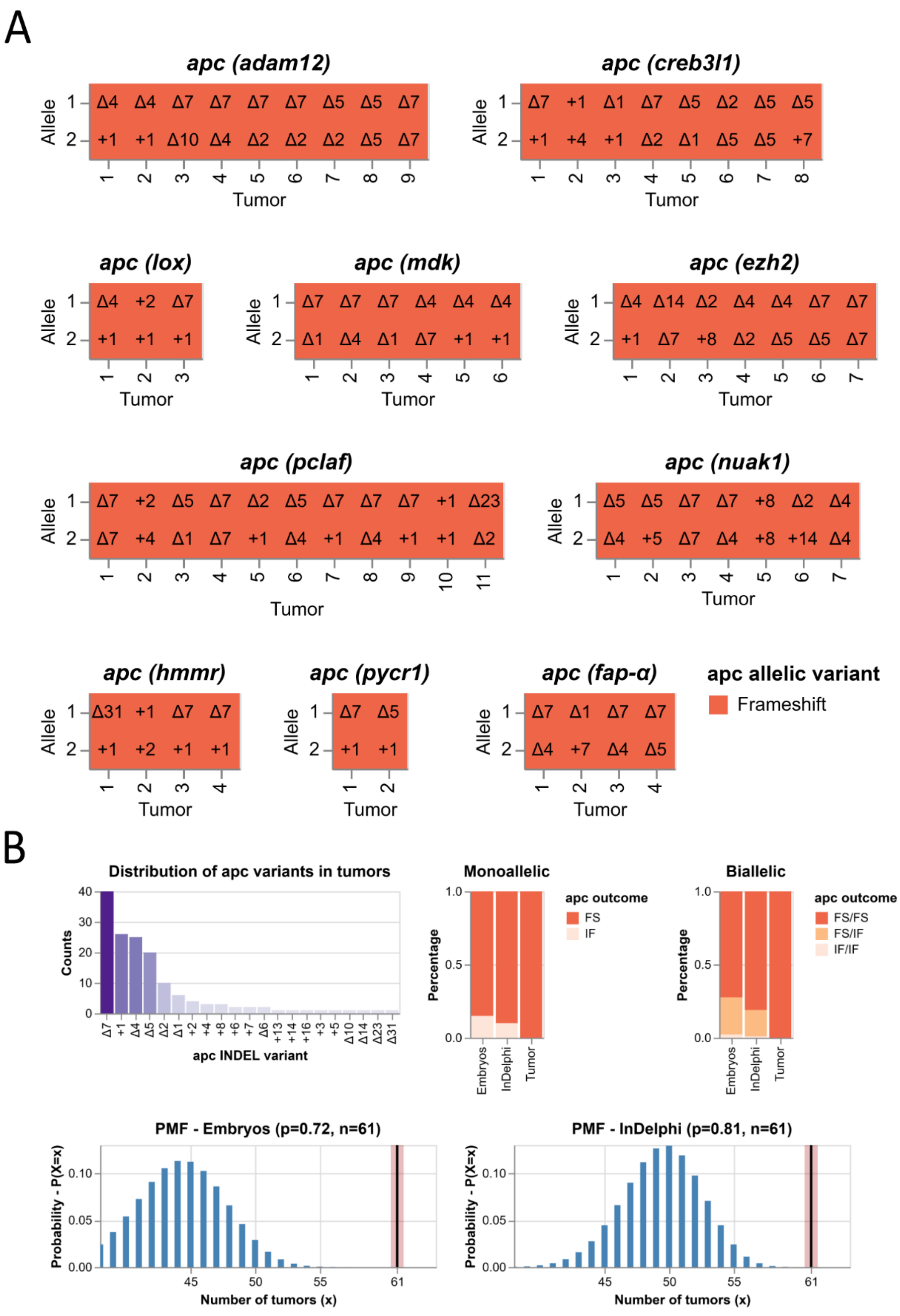


**Figure S1: The mutational spectrum of *apc* in desmoid tumors reveals positive selection for biallelic frameshift mutations. (A)** *Xenopus tropicalis* embryos are co-targeted at *apc*

and respectively one suspected genetic dependency (*adam12*, *creb3l1*, *lox*, *mdk*, *ezh2*, *pclaf*, *nuak1*, *hmmr1*, *pycr1* or *fap-α*). Desmoid tumors were dissected from post-metamorphic animals (aged three months) and both CRISPR/Cas9 target sites were subjected to targeted amplicon sequencing to determine gene editing outcomes. Shown here are the editing outcomes at the *apc* locus. **(B)** The probability to sample a biallelic frameshift mutation (right) in a desmoid tumor is directly related to the probability on monoallelic frameshift editing outcomes (left). *E.g.* upon an *apc* gRNA-specific frameshift frequency of 85%, the probability of a single desmoid tumor to be biallelic frameshift mutant is 72% (0.85*0.85). **(C)** (left) Given the *apc* editing outcomes sampled in embryos, *i.e.* in absence of selective pressure, the probability of a single desmoid tumor to have biallelic frameshift editing outcomes is 72%. Therefore, the probability of sampling 94% (61/61 tumors) with biallelic frameshift allelic status is very unlikely (probability < 0.001) according to binomial theory. (right) Similarly, given *apc* editing outcomes predicted by InDelphi, the probability of a single desmoid tumor to have biallelic frameshift editing outcomes is 81%. Therefore, the probability of sampling 94% (61/61 tumors) with biallelic frameshift allelic status is very unlikely (probability < 0.001) according to binomial theory.

**Supplementary Figure 2:**

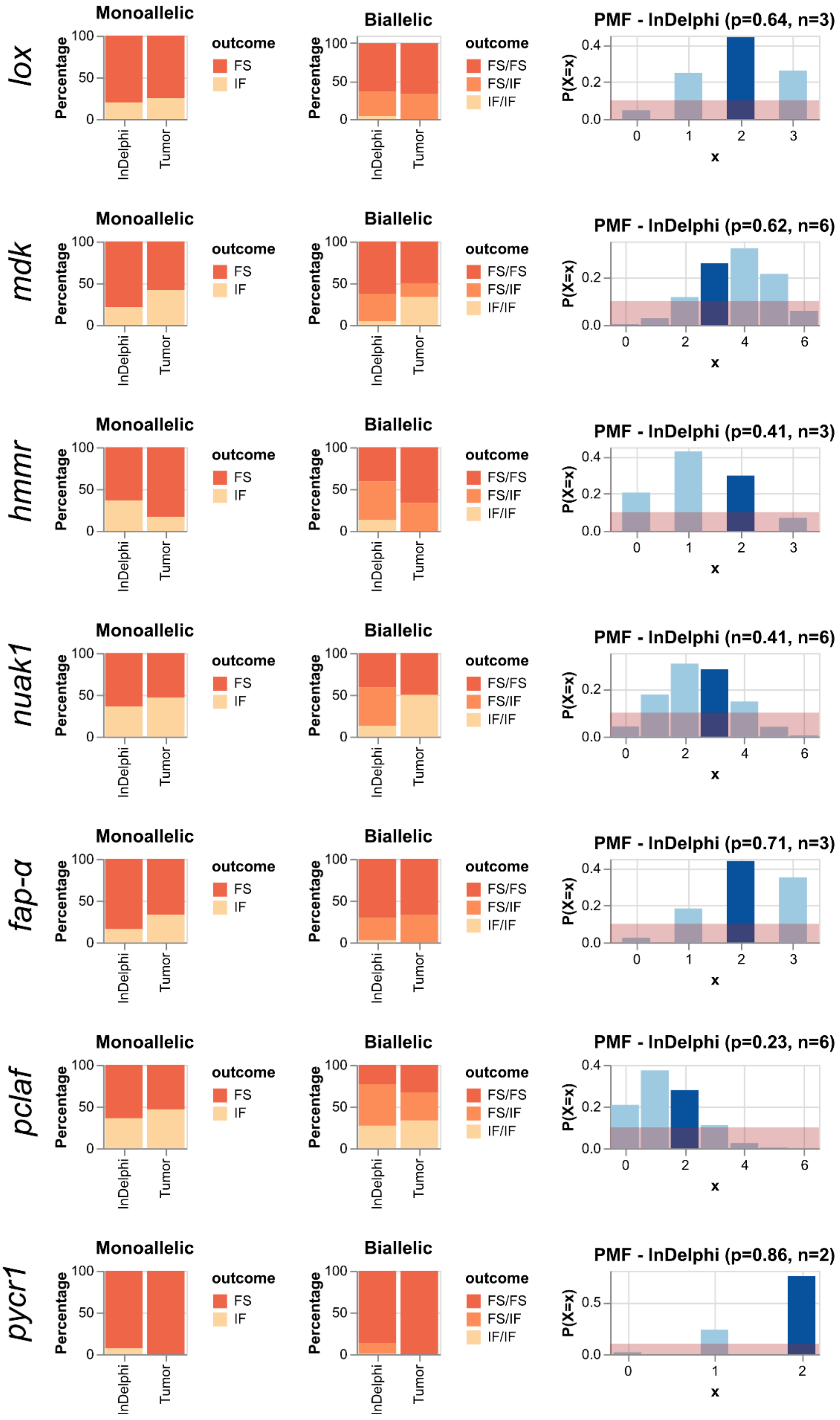


**Figure S2: No selection towards specific patterns of gene editing outcomes at suspected dependency target sites in *apc* CRISPR/Cas9-induced desmoid tumors for *lox*, *mdk*, *hmmr*, *nuak1*, *fap-α*, *pclaf*, *pycr1*.** Gene editing outcomes at suspected dependencies demonstrate, for each gene, at least two tumors developing with biallelic frameshift mutations. Gene editing outcomes in desmoid tumors are in line, and probable according to binomial theory, with predicted gene editing outcomes as determined by the InDelphi algorithm. *E.g* for *lox*, given the editing outcomes as predicted by InDelphi, the probability of a single biallelic mutant desmoid tumor to have biallelic frameshift editing outcomes is 64%. The probability of sampling 66% (2/3 tumors) with biallelic frameshift *lox* status is likely (probability is 44%). Red demarcation represents a 10% probability interval.

**Supplementary Figure 3:**

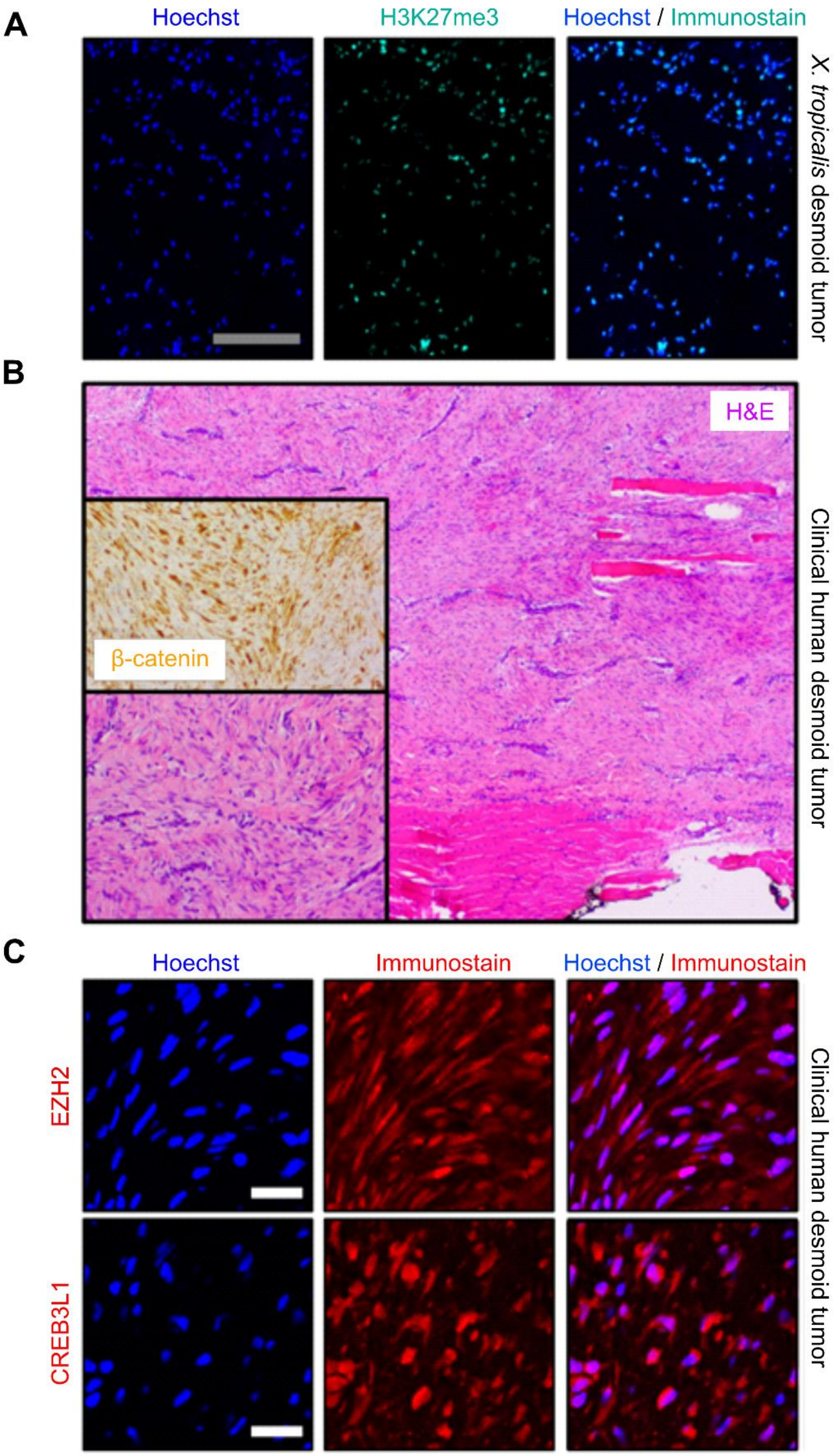


**Figure S3: H3K27me3 immunoreactivity in *X. tropicalis* desmoid tumors. Further, immunostaining reveals CREB3L1 and EZH2 expression in clinical human desmoid tumors.** **(A)** *Xenopus tropicalis* desmoid tumor cells demonstrate immunoreactivity for H3K27Me3. Grey scale bar is 200 µm. **(B)** Representative photomicrograph for three case studies of desmoid tumors employed for immunostainings in this study. First, irregular infiltration of DT cells into adjacent skeletal muscle could be noted. (Inset) Higher resolution investigation revealed proliferation of elongated, slender, spindle-shaped cells of uniform appearance, set in a collagenous stroma containing prominent blood vessels. The cells lack nuclear hyperchromasia or cytological atypia and are arranged in long sweeping bundles. (Inset; β-catenin) The spindle cells show cytoplasmatic and nuclear positivity for β-catenin. Taken together, these histopathological hallmarks were compatible with the diagnosis of DT in all three case studies. Immunostaining on these clinical samples revealed both nuclear and cytoplasmatic reactivity with anti-EZH2 **(C-top)** and anti-CREB3L1 **(C-bottom)** antibodies. Please note that CREB3l1 is normally inserted into ER membranes, with the N-terminal DNA-binding and transcription activation domains oriented toward the cytosolic face of the membrane. Upon activation and cleavage in the Golgi, the CREB3L1 N-terminal fragment will translocate to the nucleus. The CREB3L1 N-terminal-reactive antibody (AF4080; Rndsystems) demonstrates both nuclear and cytoplasmatic immunoreactivity indicative for a role of CREB3L1 as an active transcription factor in desmoid tumors. Shown immunofluorescence is representative for all three desmoid tumor case studies. White scale bar is 20 µm.

#### Supplementary Figure 4:

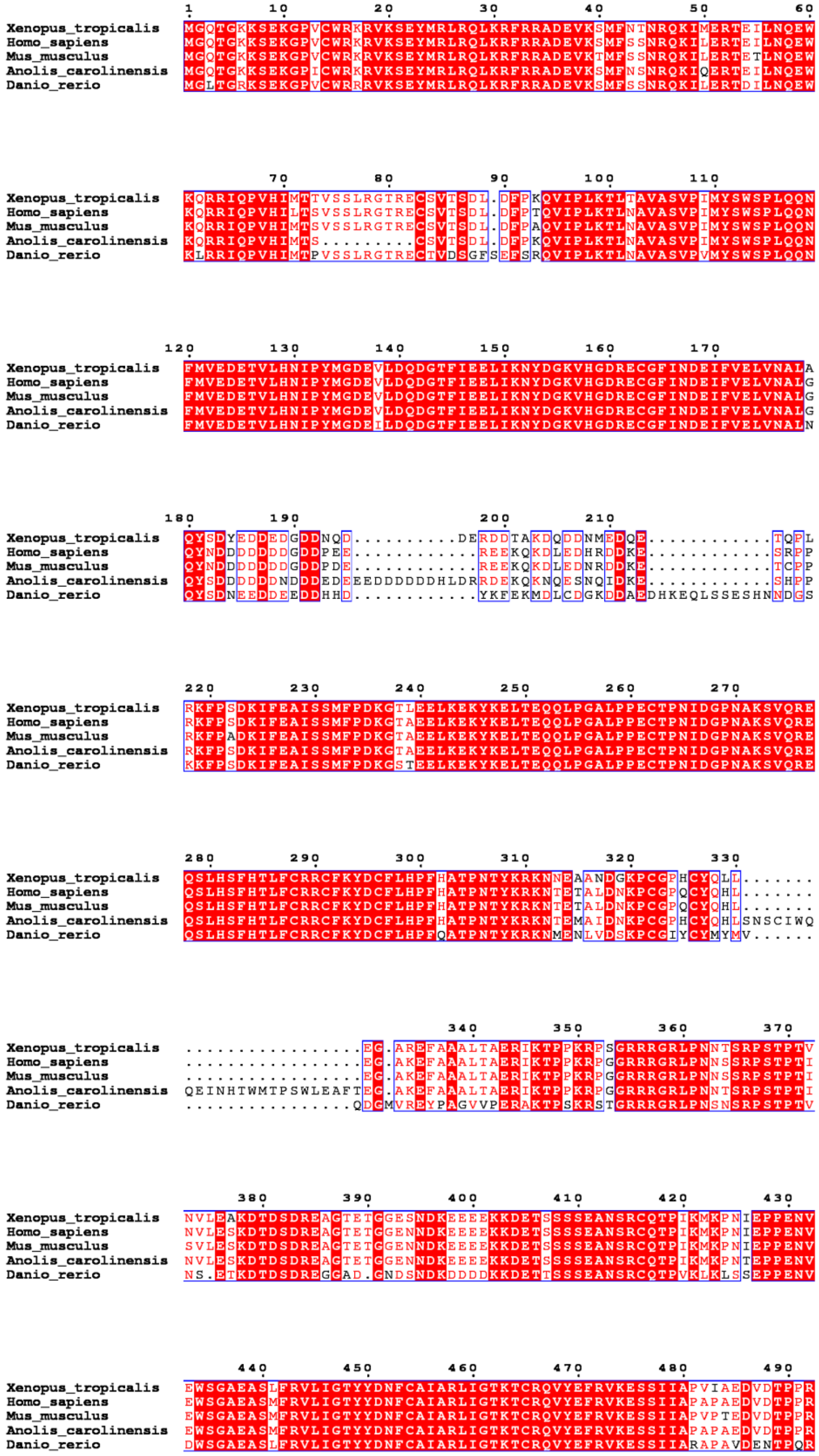

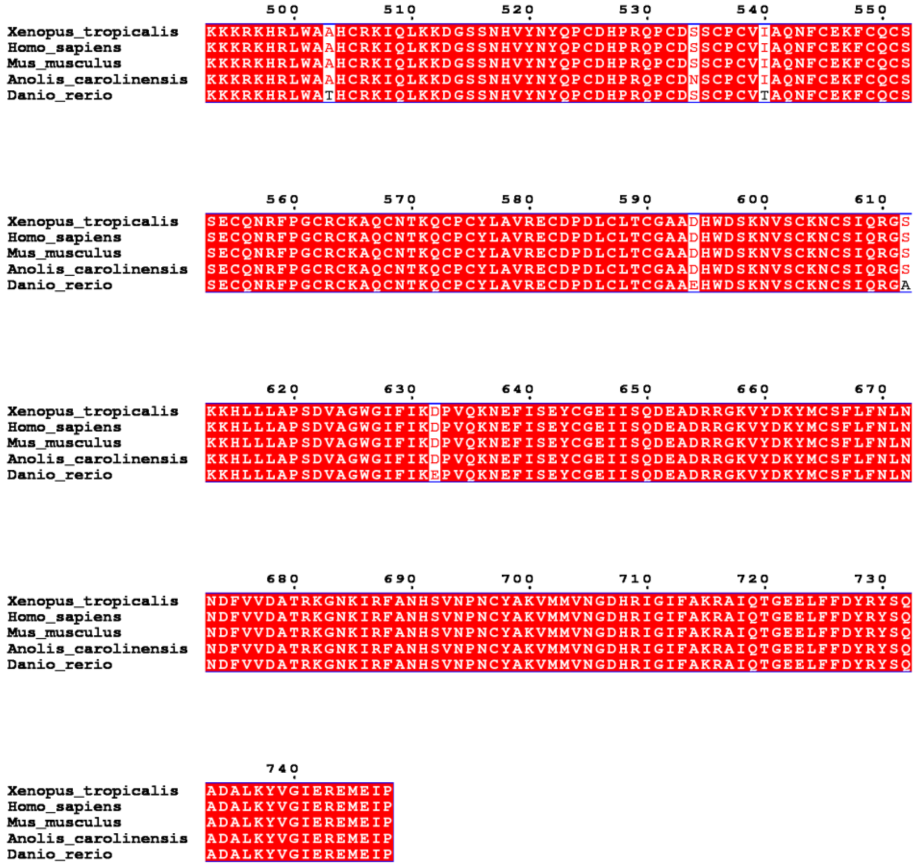


**Fig. S4: Cross-species multiple sequence alignment for EZH2 protein.** The structural template shown is based on pdb entry 5IJ7[29]. Ezh2 from *Xenopus tropicalis* shares 96% sequence identity with the *Ac*EZH2 used to assemble the structural template in Fig 4B. Further are the regions containing mutations 100% conserved between human EZH2, *X. tropicalis* Ezh2 and *Ac*EZH2 (See alignment). Followed UniProtKB entries were used: Q61188 (EZH2_MOUSE), Q15910 (EZH2_HUMAN), R4GB81 (R4GB81_ANOCA), Q08BS4 (EZH2_DANRE), Q28D84 (EZH2_XENTR).

**Supplementary Figure 5:**

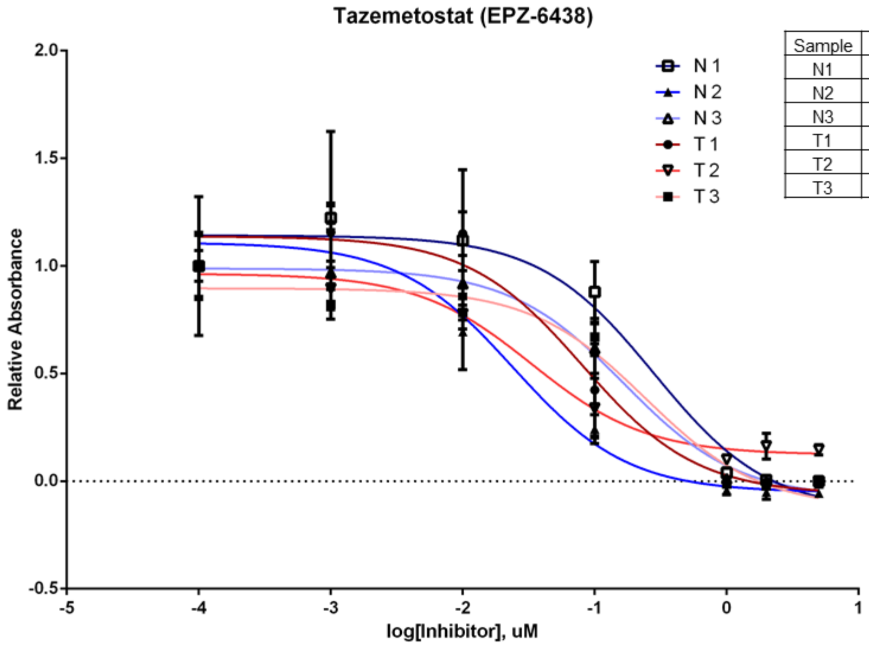


| Sample | IC50 (uM) | 95% CI |
|---|---|---|
| N1 | 0.2827 | 0.1178 to 0.6783 |
| N2 | 0.02411 | 0.009382 to 0.06194 |
| N3 | 0.1665 | 0.1152 to 0.2407 |
| T1 | 0.08091 | 0.03635 to 0.1801 |
| T2 | 0.03352 | 0.01743 to 0.06447 |
| T3 | 0.243 | 0.1225 to 0.4822 |

**Fig. S5: Proliferation characteristics of three desmoid tumor cell lines (T1-3) and three matched normal fibroblast cultures (N1-3) are similar after tazemetostat treatment.** Dose-response curve for two-week treatments with Tazemetostat at the following concentrations: 5 µM, 2 µM, 1 µM, 0.1 µM, 0.01 µM, 0.001 µM, 0.0001 µM. Shown is absorbance at 370 nm relative to BrdU incorporation added 18h before experiment end-point. Non-linear regression of “Relative Absorbance vs. log[Drug]” and IC50 calculations were conducted using GraphPad Prism. Error bars are standard deviation. 95% confidence intervals of IC50 are reported in the table.

**Supplementary Figure 6:**

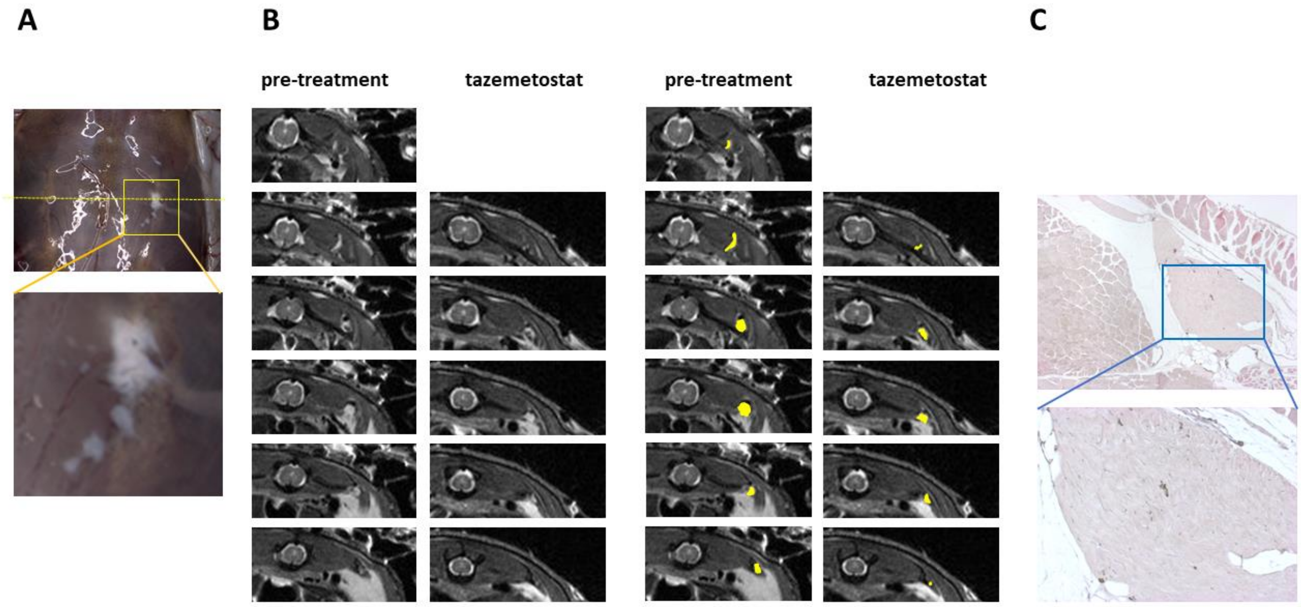


**Figure S6. Application of MRI to follow drug response.** (**A**) Desmoid tumors in the dorsal muscle of a 4-year old *apc*$^{MCR\text{-}\Delta 1/+}$ animal exposed for 4 weeks to 10 µM Tazemetostat. (**B**) Sequential transversal MRI scans showing a desmoid tumor (yellow outline) in between the dorsal muscles before (left) and after (right) Tazemetostat treatment. (**C**) H&E stained histological section of desmoid tumor visible in the MRI scan.

**Supplementary Figure 7:**

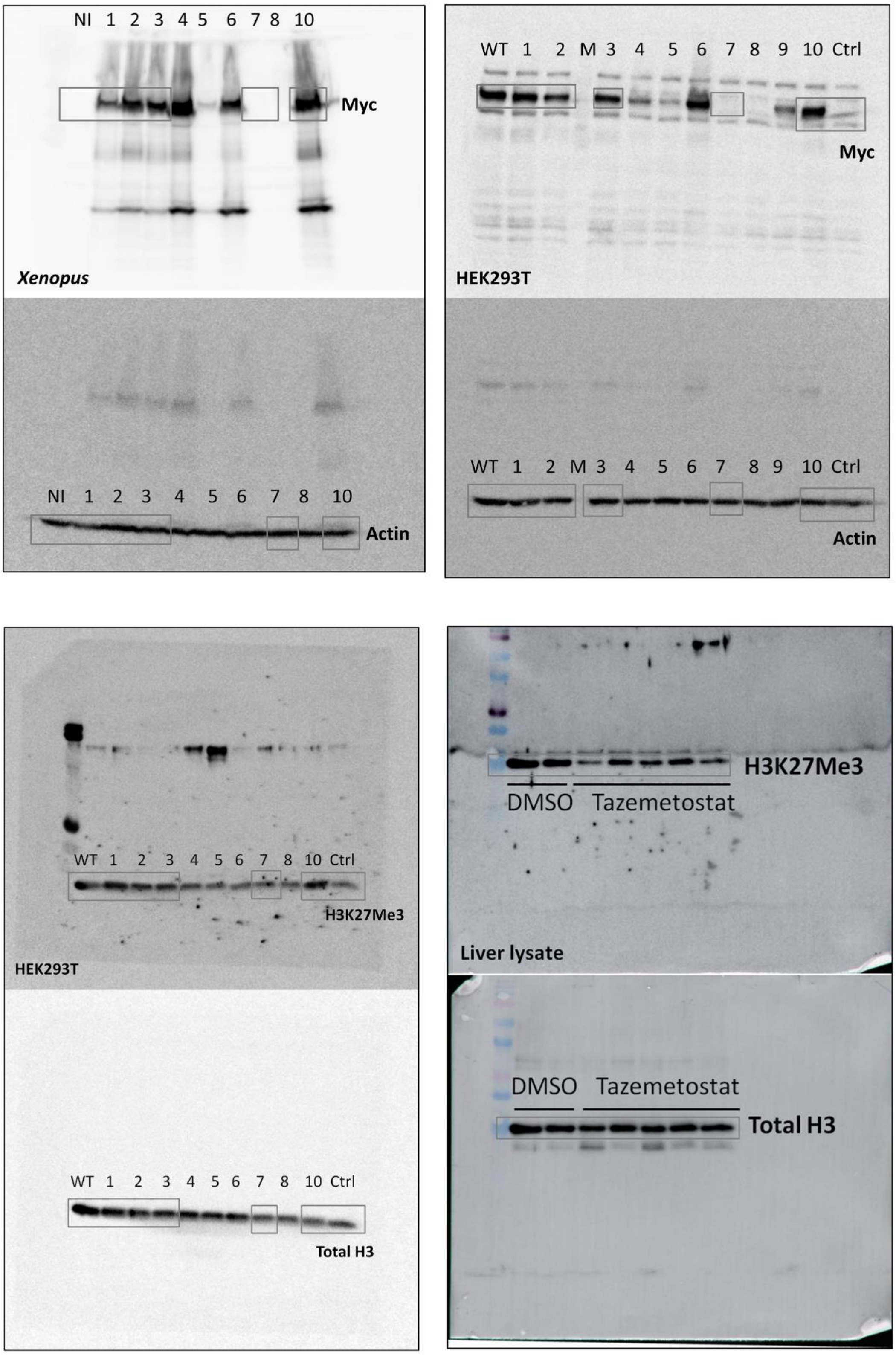


**Figure S5. Uncropped western blots.** Following legend was used. Parts of these blots used in main Fig. 4C and Fig. 5B are demarcated by grey boxes.

(NI) not injected
**(1) NM_00101793.2:c.2163_2180del; Ezh2(685-KIRFANHSVNPNCYAK-700); deletion18, variant1**
**(2) NM_001017293.2:c.2165_2182del; Ezh2(685-KIRFANHSVNPNCYAK-700); deletion18, variant2**
**(3) NM_001017293.2:c.2171_2179del; Ezh2(685-KIRFANHSVNPNCYAK-700); deletion9**
(4)NM_001017293.2:c.143_178del;
(5) NM_001017293.2:c.127_147del;
(6) NM_001017293.2:c.144_149del;
(7) NM_001017293.2:c.2068_2074del; Ezh2(p.Lys658fs)
(8) NM_001017293.2:c.2068del;
(9) NM_001017293.2:c.2170_2171insAAACAA;
(10) NM_001017293.2:c.2127A>T,
(ctrl) pMAX-GFP

### Supplementary Tables (caption Excel files)

**Supplementary Table S1:** Genotyping by PCR amplification, sequencing (MiSeq) and BATCH-GE analysis

**Supplementary Table S2:** Results of differential expression analysis and gene set enrichment analysis of the genes upregulated in desmoid tumors compared to other fibrotic lesions

**Supplementary Table S3:** Compiled list of potential desmoid tumor genetic dependencies

**Supplementary Table S4:** CRISPR-SID - deep amplicon sequencing of candidate genetic dependencies in dissected desmoid tumors

**Supplementary Table S5:** CRISPR-SID analysis of EZH2 and CREB3L1 as genetic dependencies

**Supplementary Table S6:** Statistical Analyses

**Supplementary Table S7:** Sequences guide RNAs and oligos and CRISPR/Cas9 injection setups

> **Table S7A**: guide RNA genomic target sites and oligos used for cloning-free generation
>
> **Table S7B**: Genotyping primers for CRISPR/Cas9 target sites
>
> **Table S7C**: CRISPR/Cas9 injection injection set-ups
>
> **Table S7D**: List of primers and sequences used for qRT-PCR experiments
